# Whole-Brain fMRI Functional Connectivity Signatures Predict Sustained Emotional Experience in Naturalistic Contexts

**DOI:** 10.1101/2022.11.08.515743

**Authors:** Shuyue Xu, Zhiguo Zhang, Linling Li, Yongjie Zhou, Danyi Lin, Li Zhang, Gan Huang, Xiqin Liu, Benjamin Becker, Zhen Liang

**Affiliations:** School of Biomedical Engineering, Health Science Center, Shenzhen University, Shenzhen 518060, China; Guangdong Provincial Key Laboratory of Biomedical Measurements and Ultrasound Imaging, Shenzhen 518060, China; Institute of Computing and Intelligence, Harbin Institute of Technology, Shenzhen, China; Peng Cheng Laboratory, Shenzhen 518055, China; Marshall Laboratory of Biomedical Engineering, Shenzhen 518060, China; Department of Psychiatric Rehabilitation, Shenzhen Kangning Hospital, Shenzhen, China; Center of Psychosomatic Medicine, Sichuan Provincial Center for Mental Health, Sichuan Provincial People’s Hospital, MOE Key Laboratory for Neuroinformation, University of Electronic Science and Technology of China, Chengdu 611731, China

**Keywords:** naturalistic stimulation, fMRI, functional connectivity, emotion prediction, movie clips

## Abstract

Determining and decoding emotional brain processes under ecologically valid conditions remains a key challenge in affective neuroscience. The current functional magnetic resonance imaging (fMRI) based emotion decoding studies are mainly based on brief and isolated episodes of emotion induction, while sustained emotional experience in naturalistic environments that mirror daily life experiences are scarce. Here we use 10-minute movie clips as ecologically valid emotion-evoking procedures in n=52 individuals to explore emotion-specific fMRI functional connectivity (FC) profiles on the whole-brain level at high spatial resolution (400 atlas based parcels). Employing machine-learning based decoding and cross validation procedures allowed to develop predictive FC profiles that can accurately distinguish sustained happiness and sadness and that generalize across movies and subjects. Both functional brain network-based and subnetwork-based emotion prediction results suggest that emotion manifests as distributed representation of multiple networks, rather than a single functional network or subnetwork. Further, the results show that the Visual Network (VN) and Default Mode Network (DMN) associated functional networks, especially VN-DMN, exhibit a strong contribution to emotion prediction. To further estimate the cumulative effect of naturalistic long-term movie-based video-evoking emotions, we divide the 10-min episode into three stages: early stimulation (1 ~ 200 s), middle stimulation (201 ~ 400 s), and late stimulation (401 ~ 600 s) and examine the emotion prediction performance at different stimulation stages. We found that the late stimulation has a stronger predictive ability (accuracy=85.32%, F1-score=85.62%) compared to early and middle stimulation stages, implying that continuous exposure to emotional stimulation can lead to more intense emotions and further enhance emotion-specific distinguishable representations. The present work demonstrates that sustained sadness and happiness under naturalistic conditions are presented in emotion-specific network profiles and these expressions may play different roles in the generation and modulation of emotions. These findings elucidate the importance of network level adaptations for sustained emotional experiences during naturalistic contexts and open new venues for imaging network level contributions under naturalistic conditions.

## Introduction

Human emotion represents a dynamic process involving different levels of processing and integration (Cowen and Keltner, 2017; Horikawa et al., 2020). Determining the specific neurophysiological basis of emotions and their distinct neural representations can facilitate discriminating specific emotional states from other mental processes (Putkinen et al., 2021; Saarimäki et al., 2022; Vytal and Hamann, 2010; Zhou et al., 2020; Zhou et al., 2021) and help evaluate emotion-specific dysregulations and corresponding treatment approaches (Reddan et al., 2018; Xu et al., 2022; Zhang et al., 2022). To capture the complex and dynamic nature of emotional processes, an increasing number of functional Magnetic Resonance Imaging (fMRI) studies examined cortical and subcortical responses during naturalistic contexts such as narratives or movies (Jääskeläinen et al., 2021; Putkinen et al., 2021; Saarimäki et al., 2022). Recent neuroimaging studies have shown that distributed activity and connectivity patterns underly emotion perception and processing (Baucom et al., 2012; Kassam et al., 2013; Saarimäki et al., 2016; Zhou et al., 2020; Zhou et al., 2021). However, the existing fMRI-based emotion studies seldom explore how emotions are represented under sustained and dynamic emotional engagement which better mirrors emotional processes in everyday life. To better understand emotion-related brain states under a naturalistic condition, we introduce a long-term naturalistic continuous emotion-evoking paradigm based on a number of movie clips. We chose comparably long-term movie clips to establish a naturalistic paradigm, which allows to neurofunctionally map sustained emotional experience in real-world environments (Sonkusare et al., 2019). This naturalistic fMRI approach could offer a more powerful and reliable strategy to capture the complexity and variation of emotions across subjects and allow to generate ecologically valid brain signatures of emotional processes (Eickhoff et al., 2020).

Recent network-level based perspective on emotional brain processes suggest that dynamic interactions between brain regions play a significant role in emotional experience and regulation. To examine complex changes within integrative brain networks and interpret large-scale neuronal communication, functional connectivity (FC) has gained increasing interest. This approach provides a powerful tool to examine connectivity changes and complex integrative brain networks and to further determine the role of large-scale networks in emotional states, as has been demonstrated robust across individuals and task paradigms (Betti et al., 2013; Vanderwal et al., 2017; Wang et al., 2017b). FC reflects functional interaction between anatomically separated brain regions and is thought to reflect the temporal dependency of brain regions in terms of neural activation patterns. FC has been increasingly suggested to represent a robust biomarker for mental processes and their dysregulation, including emotion (Magalhães et al., 2021; Putkinen et al., 2021; Saarimäki et al., 2022; Zhuang et al., 2021), cognition (Cohen, 2018; Ptak et al., 2020; Zimmermann et al., 2018a), developmental changes (Ciarrusta et al., 2020; Liu et al., 2021; Teeuw et al., 2019), and disorders (Du et al., 2018; Xu et al., 2021; Zheng et al., 2018; Zhou et al., 2018; Zimmermann et al., 2018b). It has been further demonstrated the feasibility of analyzing FC at the level of concordant patterns of temporal variations under the mental states during rest and task (Cohen and D’Esposito, 2016). Recently, a number of FC-based emotion-related brain network studies have been conducted (Pessoa, 2017, 2018; Putkinen et al., 2021; Saarimäki et al., 2022). For example, Putkinen et al. examined the emotion-related functional neural basis of four separate emotions evoked by music (happiness, sadness, fear, and tender) and observed that brain activity patterns in the auditory and primary motor cortices correlated with the respective emotional states (Putkinen et al., 2021). Saarimäki et al. introduced multivariate pattern analysis to develop a cross-subject emotion recognition approach that is based on whole-brain FC profiles. The study collected brain activity in 16 subjects during fMRI while the subjects were presented with 1-minute emotional audio narratives from six emotion categories (anger, fear, disgust, happiness, sadness, and surprise) (Saarimäki et al., 2022). The results showed that the most accurate emotion prediction could be obtained from the Default Mode Network (DMN), indicating an important contribution of the DMN in emotional processing under naturalistic conditions. Collectively, these studies have demonstrated the feasibility to detect emotional changes via FC patterns in naturalistic environments. The high discriminative power of FCs has moreover been demonstrated in the classification of emotion-related diseases (Wang et al., 2020; Zeng et al., 2012; Zeng et al., 2014). However, most previous emotion decoding studies relied on a few minutes of fMRI data, such as 45-second music (Putkinen et al., 2021) or 1-minute narratives (Saarimäki et al., 2022). Although these studies demonstrated the feasibility to decode specific emotions from FC patterns, the performance varied between emotional categories, in particular the specificity and accuracy for sadness remained limited. For instance, despite a high accuracy obtained for most emotions in Saarimäki et al.’s study (Saarimäki et al., 2022), it revealed a classification accuracy of sadness was close to chance level (18% accuracy). One possible reason might be the different time frames of emotional experiences and thus the difficulty in robustly evoking strong and engaging feelings of sadness with experimental stimuli as short as one minute. In contrast to emotions such as fear, surprise, or general negative affect which can be reliably induced by short and sparse stimuli (Čeko et al., 2022; Xin et al., 2020; Zhou et al., 2021), a strong subjective experience of sadness may require a longer timeframe and more contextual information. To this end, longer immersive experimental stimuli may facilitate the induction of robust emotional experiences and allow more robust decoding (Waugh and Kuppens, 2021; Waugh et al., 2012).

On the other hand, recent studies using naturalistic movie-watching paradigms (Demirtaş et al., 2019; Gilson et al., 2018; Kim et al., 2018; Ren et al., 2018; Sonkusare et al., 2019) have shown that the estimated FC during movie-watching exhibits a high test-retest reliability and may allow to capture brain function under naturalistic contexts matching real-life processing (Di and Biswal, 2020; Di et al., 2021; Wang et al., 2017b). In the context of emotion research, naturalistic movies in combination with fMRI could offer a more powerful and reliable tool for capturing the complexity and variation of emotions across subjects and under ecologically more valid conditions of emotional processes (Eickhoff et al., 2020). However, to our knowledge, a whole-brain network analysis under naturalistic long movie stimulation (e.g. 10-minute) for emotion prediction has not been conducted. In turn, the current evidence for emotion-related cortical and subcortical engagement in naturalistic contexts - especially for long and dynamic emotional experiences remains elusive (Kragel and LaBar, 2014; Lindquist and Barrett, 2012; Lindquist et al., 2012). To this end, the present study aimed to address the following open questions:

1. Which brain networks and connections exhibit emotion-specific contributions to the sustained emotional state on the whole-brain level?
2. To which extent do the network level profiles vary over time during the sustained emotion induction procedure?

To address these questions, we designed a naturalistic emotion induction paradigm including 12 different movie clips with a duration of 10 minutes and simultaneously recorded fMRI data in n = 52 healthy participants. We primarily focus on two distinct basic emotions, happiness and sadness, which cover the positive and negative valence dimension. To capture the hemodynamic brain changes during the movie clip on the whole-brain functional connectivity network level, a high spatial resolution parcellation with 400 regions-of-interest from the Schaefer 400 atlas (Schaefer et al., 2018) was adopted to generate emotion-related network level changes. Next, a cross-subject cross-episode emotion prediction model based on whole-brain-level FC analysis was developed and evaluated using a cross-subject leave-one-subject-out cross-validation. This data-driven analysis using the whole-brain FC patterns for the prediction of two distinct basic emotions (happiness and sadness) allowed validation of the decoding models. The current study further allowed us to better examine the communications between different key brain regions during emotional processing under a naturalistic stimulation paradigm with movie clips. The decodability and generalizability of different functional networks on emotions across different subjects and movie clips were examined while the contributions of the brain areas during emotion evoking under naturalistic movie clips could be determined and important whole-brain connectivity patterns were evaluated.

## Experimental Materials

### Subjects

A total of 52 healthy right-handed subjects (male/female: 26/26; age: 19 to 28 years old, 23.52±2.05; with normal or corrected-to-normal vision) from Shenzhen University were recruited to participate in the experiment. Exclusion criteria included neurological or psychiatric diagnosis, heavy alcohol consumption within the past 6 months, cardiovascular disease, and severe visual impairment. All the subjects signed the informed consent before starting the experiment, and the experiment was approved by the Ethics Committee of the Health Science Center, Shenzhen University. The experiment was in line with the latest Version of the Declaration of Helsinki. After quality control, data from one subject (female; age: 19 years old) was excluded due to high head motion.

### Stimuli

Twelve ten-minute complex and naturalistic movie clips with strong and reliable emotion eliciting effects were selected from 12 different natural-colored movies. The twelve 10-minute movie clip candidates include 6 happy stimuli (positive emotion) and 6 sad stimuli (negative emotion). There is no content overlap between the selected movie clips. The 6 happy stimuli were from the movies of “Mr. Popper’s Penguins”, “Ted”, “The Onion Movie”, “Liar Liar”, “A Thousand Words”, and “Absolutely Anything”, and the 6 sad stimuli were from “Miracle In Cell No.7”, “Prayers For Bobby”, “The Classic”, “Grave Of The Fireflies”, “Only The Brave”, and “The Last Train”. More details about the stimuli selection are reported in Appendix I of the Supplementary Materials.

### Experimental Paradigm

The experimental paradigm is shown in Fig. 1. For each subject, the fMRI experiment under naturalistic stimulation included a total of 6 episodes (corresponding to 6 movie clips). The 6 movie clips were randomly selected from the original twelve 10-minute movie clip candidates, including 3 happy stimuli and 3 sad stimuli. The average selection rate of each stimulus is 8.33±1.28%. The presentation order was randomized to counterbalance the order effect. Similar to previous studies (Horikawa et al., 2020; Saarimäki et al., 2016), the movie clips were presented without sound. For a single episode, it comprised a 30-second baseline (subjects looked at the white cross shown in the center of a black screen to clear their mind), 10-minute movie playing (subjects passively viewed the movie clips with full engagement), and subjective feedback (subjects rated their emotional experience evoked by the presented movie clips). At the end of one episode, the subjects could take a self-paced break. Throughout the whole procedure, the subjects were requested to remain still.

**Figure 1.**
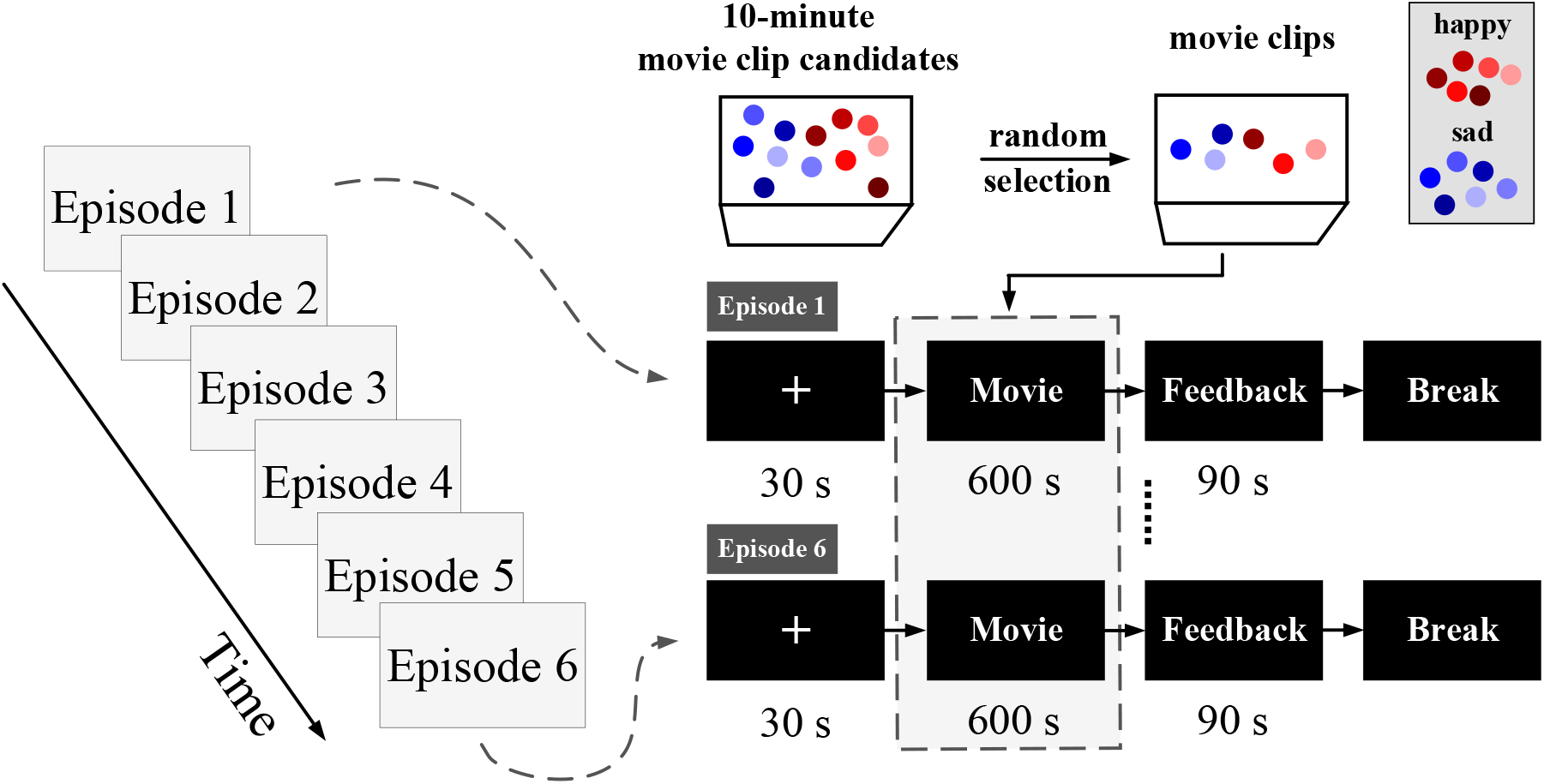
The experimental paradigm. For each subject, the experiment includes 6 episodes with a randomized playing sequence of the selected movie clips. Here, the 6 movie clips, including 3 happy movie clips and 3 sad movie clips, are randomly selected from a total of 12 10-minute movie clip candidates.

Brain images were recorded using a Siemens Prisma 3 Tesla MRI scanner with a 64-channel head coil. High-resolution T1-weighted structural images covering the entire brain were acquired using a magnetization-prepared rapid acquisition gradient echo (MPRAGE) sequence with the parameters as: voxel resolution=1×1×1 mm^3^, repetition time (TR)=2300 ms, echo time (TE)=2.26 ms, field of view (FOV)=256×232 mm^2^, flip angle (FA)=8°. During the experimental task, the functional images were acquired using a single gradient echo-planar imaging (EPI) sequence, with the parameters as: voxel resolution=2×2×2 mm^3^, TR=1000ms, TE=30ms, FOV=192× 192mm^2^, FA=90°. Each volume of EPI functional images consisted of 65 slices. During scanning, all subjects were instructed to remain awake, keep eyes open, and be in full engagement with the presented movie clips. All subjects completed six functional scans according to the six 10-minute natural movie clips presented via E-Prime version 3.0. Movie clips were counter-projected on a screen and viewed through a mirror mounted on a head coil.

## Methods

### fMRI Preprocessing

The functional images are preprocessed using SPM (Friston, 2003) and DPARSF (Yan et al., 2016) in MATLAB. Given that the scanner included dummy scans to stabilize the magnetic field, the first five volumes of each time series are not discarded. The structural images are first stripped of the skull and segmented into gray matter (GM), white matter (WM), and cerebrospinal fluid (CSF) based on the results of the stripped skull. Then, all the EPI images of each episode are aligned with the first volume of EPI images using six head-motion parametric linear transformations, and the EPI images of each subject are aligned with the structural images. To remove the linear drift and reduce the interference of head movements and other physiological signals, head motion correction is performed where the six head movement parameters are regressed and denoised using the higher-order model (Friston 24). The EPI images are then normalized to Montreal Neurological Institute (MNI) space. A Gaussian kernel of 6 mm (FWHM) is adopted for spatial smoothing and a bandpass filter of 0.008-0.15 Hz is conducted for eliminating the low-frequency drift and high-frequency noise and improving the signal-to-noise ratio of the BOLD signal (Wang et al., 2017b).

### Segmentation of Regions of Interest

To ensure whole-brain coverage, the FC estimation is conducted using a brain parcellation atlas with 400 regions of interest (ROI) from the Schaefer 400 fMRI atlas that covers large-scale functional networks (Schaefer et al., 2018). Schaefer 400 ROIs parcellation integrates both local gradient and global similarity from rest-state and task-state FC. According to the clearly defined coordinates of the location of structural subdivisions in the whole-brain cortex, the parceled ROIs could be further categorized into 7 networks or 17 subnetworks at the coarse or fine level, respectively. As shown in Fig. 2 (a), the 7 networks include Visual Network (VN), SomatoMotor Network (SMN), Dorsal Attention Network (DAN), Ventral Attention Network (VAN), Limbic Network (LN), FrontoParietal Network (FPN), and Default Mode Network (DMN). Based on the 7 networks, the 17 subnetworks are a further subnetwork division (Fig.2 (b)), including VN-a, VN-b, SMN-a, SMN-b, DAN-a, DAN-b, VAN-a, VAN-b, LN-a, LN-b, FPN-a, FPN-b, FPN-c, DMN-a, DMN-b, DMN-c, and Temporal Parietal Network (TPN). Besides, consistent with previous studies (Luppi et al., 2022; Luppi and Stamatakis, 2021), 32 subcortical ROIs are also considered based on the recently developed Melbourne subcortical functional parcellation atlas (Tian et al., 2020), which is also obtained based on resting-state and task-state functionally connectivity and is consistent with the parcellation methodology in Schaefer 400. The 32 subcortical ROIs cover 7 subcortical regions, including hippocampus, thalamus, amygdala, caudate nucleus, caudate nucleus, putamen, and globus pallidus. All the 32 ROIs are grouped into the Subcortex Network (SN). Thus, for each episode, the corresponding fMRI data could be represented as a *r* × *t* matrix, where *r* and *t* refer to the number of ROIs and time length (*r* = 432 and *t* = 600).

**Figure 2.**
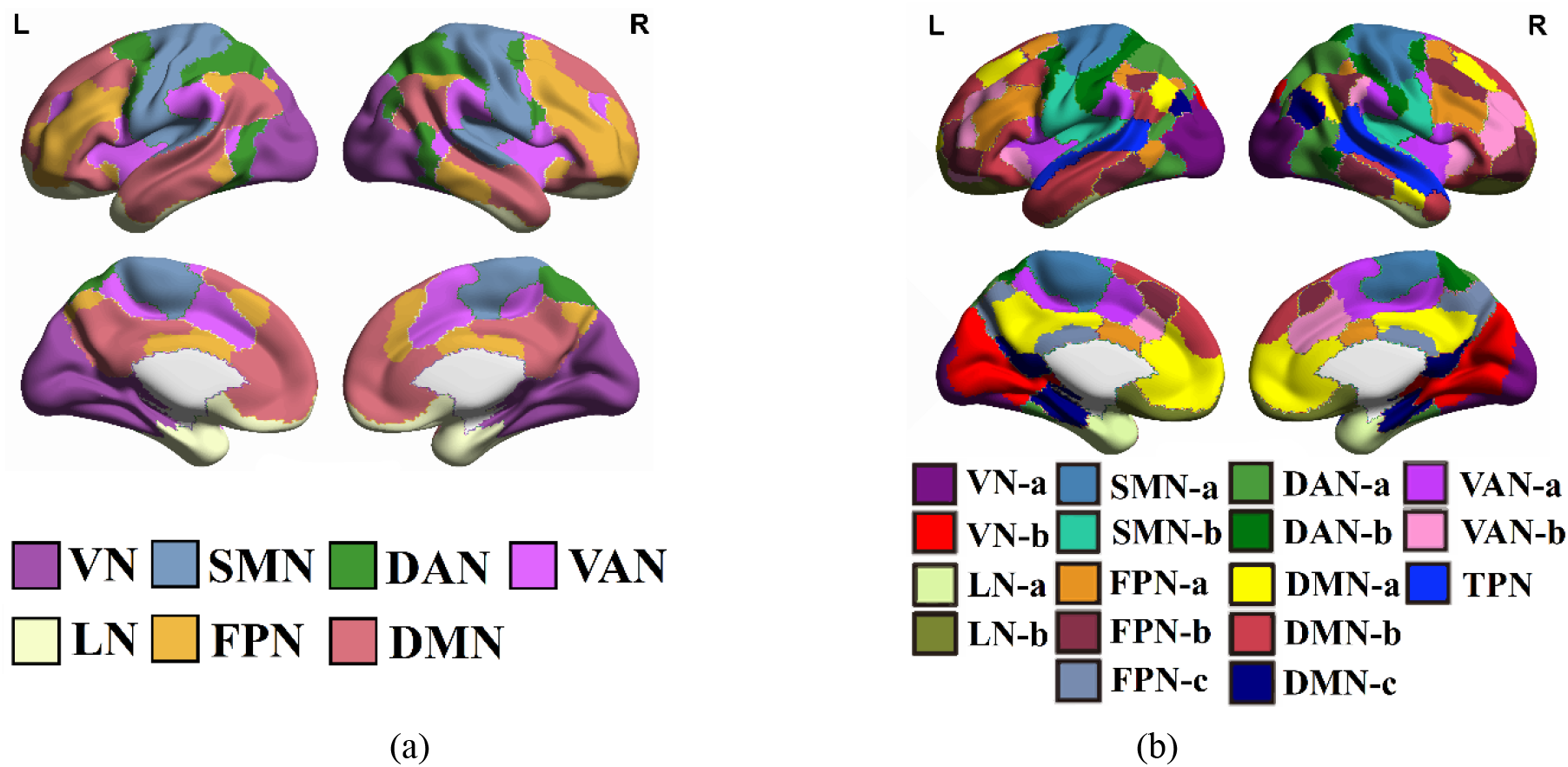
The definition of the (a) 7 networks and (b) 17 subnetworks in Schaefer 400 fMRI atlas.

### Functional Connectivity Estimation

In this study, we investigate whether the alternations of FC at different functional networks are associated with emotional states based on 432 ROIs (400 cortical regions and 32 subcortical regions). The functional signal time series of each ROI is extracted by averaging the BOLD signals of all voxels within the ROI according to the template, resulting in a BOLD time series of the size of *r* × *t* (*r* is the number of ROIs and *t* is the BOLD time series length) for each subject per episode. Here, Pearson’s correlation coefficients between the functional networks are calculated, and a FC matrix is obtained with a size of *r* × *r*. Further, the obtained FC matrix is then converted to z-values by Fisher’s z transformation for improving the normality. Each value at the normalized FC matrix indicates the strength of FC between two ROIs, which will be used as features to predict emotions in a cross-subject cross-episode manner.

### Emotion Prediction Modeling

A cross-subject cross-episode Support Vector Machine (SVM) with a linear kernel is trained to identify two distinct basic emotions (happiness and sadness). In the previous literature (Du et al., 2018), SVM is among the most commonly used classification method for cognitive and emotional brain states and achieves a stable and reliable classification performance. A total of 293 samples (subject × episodes) were used for modeling (some trials were excluded due to excessive head movements), where the number of samples corresponding to happiness and sadness is 147 and 146, respectively. In the present study, three types of episode-based emotion prediction models are built. **(1) Whole-brain-based emotion prediction modeling.** We convert the upper triangular data of the FC matrix into a (*r* × (*r* – 1))/2 by 1 feature vector to perform modeling. **(2) Network-based emotion prediction modeling.** We use the functional connection values within a network (e.g., the connection strength of ROIs in the visual network) or between two networks (e.g., the connection strength of ROIs within the visual network and the default network) as features for modeling. Since the connection matrices within networks are symmetric, only upper triangular data of the connection matrix within a network is adopted. **(3) Subnetwork-based emotion prediction modeling.** Similar to network-based modeling, we use the functional connection values within a subnetwork (e.g., the connection strength of ROIs in the central visual area) or between two subnetworks (e.g., the connection strength of ROIs within the central visual area and the peripheral visual area) as features for modeling. To verify the model effectivity and stability on cross-subject cross-episode application, the prediction performance is evaluated under a leave-one-subject-out cross-validation method which helps to verify the model generalizability on unknown subjects and episodes.

On the other hand, to estimate whether the emotion prediction performance also has an accumulative effect on the emotional experience, we examine the emotion prediction performance at different stimulation stages. Here, we divide each episode into three stimulation stages: early stimulation stage (1~200s), middle stimulation stage (201~400s), and late stimulation stage (401~600s). Each stimulation stage is a 200s duration. For each stage, a separate SVM classifier with a linear kernel is trained based on the leave-one-subject-out cross-validation protocol.

### Model Performance Evaluation

To fully evaluate the cross-subject emotion prediction performance using the extracted FC features and give an insight into the model generalizability and stability on the unknown subject(s), we conduct a strict leave-one-subject-out cross-validation. For each validation round, one subject’s all episodes are used as the test data, and the remaining subjects’ episodes are treated as the training data. The training data is used to train the prediction model, and the trained model will be then utilized on the unused test data to measure the model performance. We repeat the training validation process until each subject’s all episodes are used as the test data for once. The final model performance is an average of the obtained prediction results across all the validation rounds.

Five different evaluation metrics, classification accuracy *P_acc_*, precision *P_pre_*, sensitivity *P_sen_*, F1-Score *P_f_*, and specificity *P_spe_*, are adopted to evaluate the model performance. Suppose the correctly predicted positive and negative samples are *n_TP_* and *n_TN_* and the incorrectly predicted positive and negative samples are *n_FP_* and *n_FN_*. The classification accuracy (*P_acc_*) measures an overall classification performance, given as

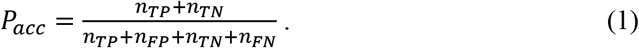

The precision (*P_pre_*) and sensitivity (*P_sen_*) are the measurement of the prediction performance on positive samples, given as

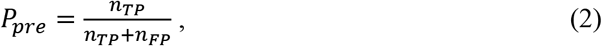

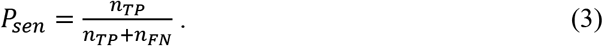

To be less susceptible to biased classification problems, we also estimate the corresponding F1-Score (*P_f_*), which is defined as

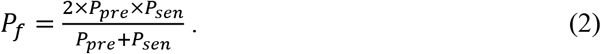

The specificity (*P_spe_*) is defined as

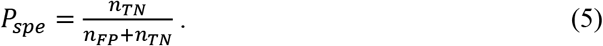

The statistical significance of all the cross-validation results is further assessed using a permutation test (Combrisson and Jerbi, 2015). To obtain a zero distribution (with a chance level of 50%: 1 divided by the number of emotion categories) for assessing prediction performance, we repeat the following procedure 1000 times to simulate the prediction probability distribution as below. (1) randomly shuffle the emotional category labels of the samples; (2) conduct leave-one-subject-out cross-validation and measure the corresponding performance by averaging the prediction accuracies in all the cross-validation rounds; (3) compare the permutation results with the truly obtained results. The p-value is calculated as the ratio of the number of accuracies in permutation results greater than the true accuracy to the total number of accuracies in permutation results. Further, to eliminate the influence of type I errors in the network-based and subnetwork-based emotion prediction modeling, all the obtained p-values are corrected using the false discovery rate (FDR) (Genovese et al., 2002) for multiple comparisons.

## Results

### Whole-brain-based emotion prediction

To estimate whether different sustained emotions would be represented in distinct FC patterns, we train a cross-subject classifier using a brain-wide FC matrix to identify the two distinct basic emotions of happiness and sadness. The average FC matrices of happiness and sadness across subjects and episodes are shown in Fig. 3 (a) and (b). The whole-brain-based emotion prediction corresponds to an accuracy of 80.55% (*P_f_*=80.55%, *P_pre_*=80.82%, *P_sen_*=80.27%, *P_spe_*=80.82%). All the results are significantly higher than the random accuracy level in the permutation test (chance level=50%; permutation test p<0.0001). The distribution of the obtained random accuracies in the permutation test (repeat 1000 times) is shown in Fig. 3 (c). The corresponding confusion matrix is shown in Fig. 3 (d), where the average prediction accuracy for happiness and sadness is 80.27% (p<0.0001) and 80.82% (p<0.0001), respectively. The results reveal that a high cross-subject cross-trial emotion prediction performance could be achieved using whole-brain functional connectivity characteristics suggesting that it is feasible to differentiate sadness and happiness based on whole-brain connectivity signatures. Next, based on the prediction performance, we determined whether different large-scale networks in terms of 8 networks and 18 subnetworks show a specific contribution to the emotion prediction.

**Figure 3.**
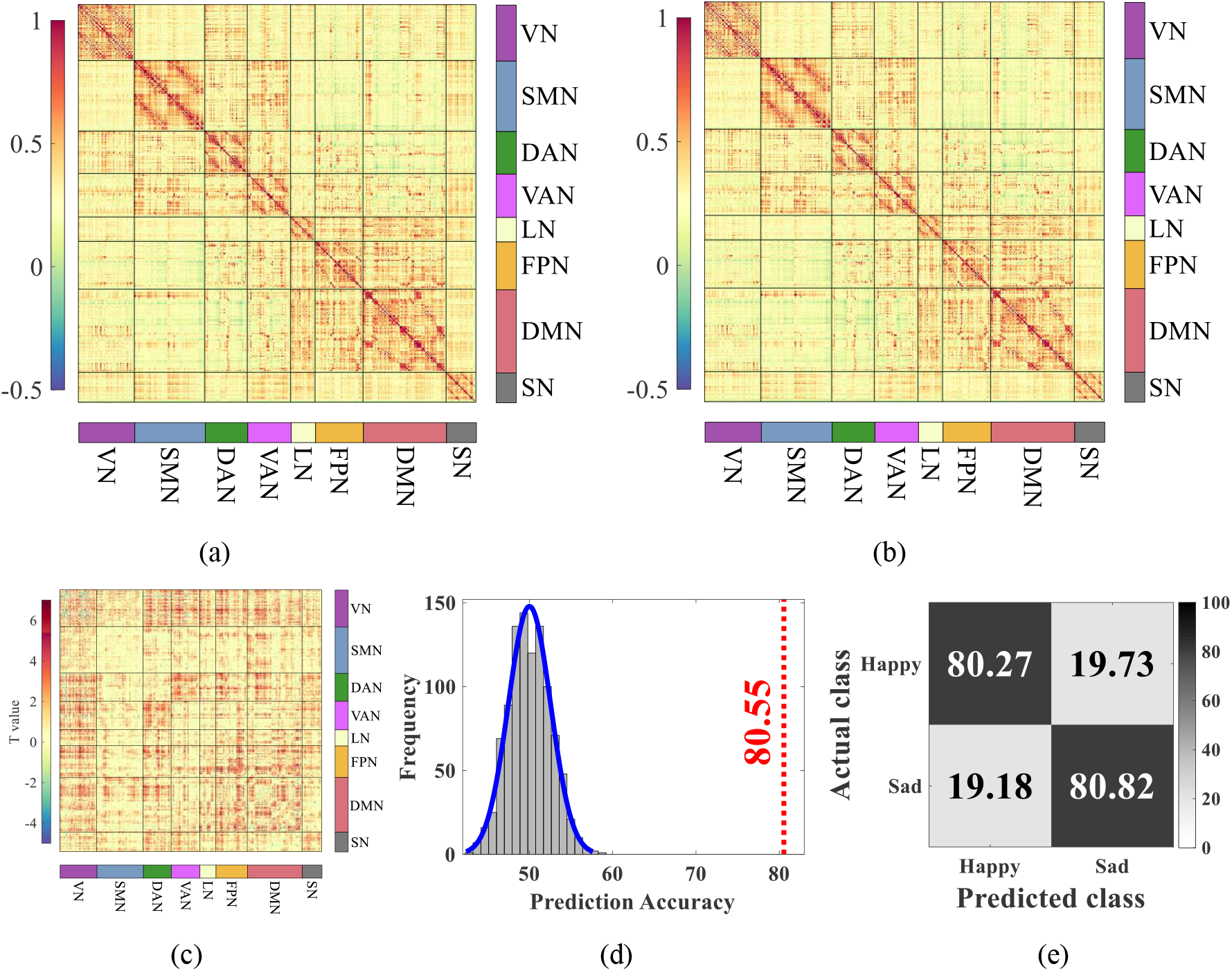
Whole-brain-based emotion prediction results. The averaged FC matrices of (a) happiness and (b) sadness. The color indicates the obtained z values after Fisher’s z transformation. (c) The statistical difference between (a) and (b). All the presented results are FDR corrected, with p<0.05. (d) The histogram distribution of the prediction accuracies is obtained in the permutation test (repeat 1000 times). The red dot line indicates the obtained whole-brain-based emotion prediction accuracy. (e) The obtained confusion matrix.

### Network-based emotion prediction

We next investigate which intra- and inter-brain functional networks are most predictive of the two distinct basic emotions (happiness and sadness). Based on the defined 8 networks, we extract the FC patterns within/between each network and build the emotion prediction model separately. The average prediction accuracy of each intra- and inter-network is reported in Fig. 4 (a). Prediction results are assessed using the permutation test, and only the classification results of both happiness and sadness are statistically greater than the random results are considered as significant (p<0.0001, FDR corrected). The obtained significance of the prediction results relative to the random level is reported in Appendix II of Supplementary Materials. The results show that the most predictive FC patterns for distinguishing happiness and sadness are mainly located in between and within network connections involving the visual network (VN) and default mode network (DMN), as shown in Fig. 4 (b). This may reflect the visual and naturalistic nature of our paradigm, which will rely on the communication between the visual system with other brain systems such as the DMN. The statistical analysis allowed us to test significant predictive networks with detailed classification performance (see Table 1). The best prediction performance is obtained when the FC between the VN and DMN is adopted (*P_acc_*=77.82%, *P_f_*=77.03%, *P_pre_*=80.15%, *P_sen_*=74.15%, *P_spe_*=81.51%; p<0.0001). The interactions between VN and DMN are in line with the economic account of large-scale brain network organization (Chen et al., 2013; Raichle, 2015; Vatansever et al., 2017; Vessel et al., 2019), which indicates the information transmissions between lower-level sensory brain areas and high-level functional brain areas. Besides, those FC patterns of VN-VAN, VN-DAN, DMN-FPN, VN, VN-SMN, DMN-LN, DMN-VAN, VN-SN, VN-LN, and DMN-SMN are also powerful to distinguish the happy and sad emotions. For all the reported results, the permutation test verifies that the probability of achieving such high prediction performance by chance is less than 0.0001 (p<0.0001, FDR corrected). The results show the functional network connections contributing to emotion prediction predominate in VN and DMN associated networks. Those network systems could be considered to involve the most informative features for emotion prediction, where a high predictive ability could be more prevalently observed.

**Table 1.** The intra- and inter-brain functional networks with the detailed prediction results. All the reported results are statistically significant (p<0.0001, FDR corrected).

| Networks | Overall Performance |  |  |  |  | Happiness | Sadness |
| --- | --- | --- | --- | --- | --- | --- | --- |
|  | Accuracy | F1-score | Precision | Sensitivity | Specificity | Accuracy | Accuracy |
| VN | 72.01% | 72.11% | 72.11% | 72.11% | 71.92% | 72.11% | 71.92% |
| **VN-DMN** | **77.82%** | **77.03%** | **80.15%** | **74.15%** | **81.51%** | **74.15%** | **81.51%** |
| VN-VAN | 75.77% | 76.09% | 75.33% | 76.87% | 74.66% | 76.87% | 74.66% |
| VN-DAN | 75.09% | 74.74% | 76.06% | 73.47% | 76.71% | 73.47% | 76.71% |
| VN-SMN | 72.01% | 72.11% | 72.11% | 72.11% | 71.92% | 72.11% | 71.92% |
| VN-LN | 69.28% | 68.53% | 70.50% | 66.67% | 71.92% | 66.67% | 71.92% |
| VN-SN | 69.62% | 69.42% | 70.14% | 68.71% | 70.55% | 68.71% | 70.55% |
| DMN-VAN | 70.31% | 70.51% | 70.27% | 70.75% | 69.86% | 70.75% | 69.86% |
| DMN-FPN | 74.40% | 75.08% | 73.38% | 76.87% | 71.92% | 76.87% | 71.92% |
| DMN-SMN | 68.60% | 69.74% | 67.52% | 72.11% | 65.07% | 72.11% | 65.07% |
| DMN-LN | 70.65% | 71.14% | 70.20% | 72.11% | 69.18% | 72.11% | 69.18% |

**Figure 4.**
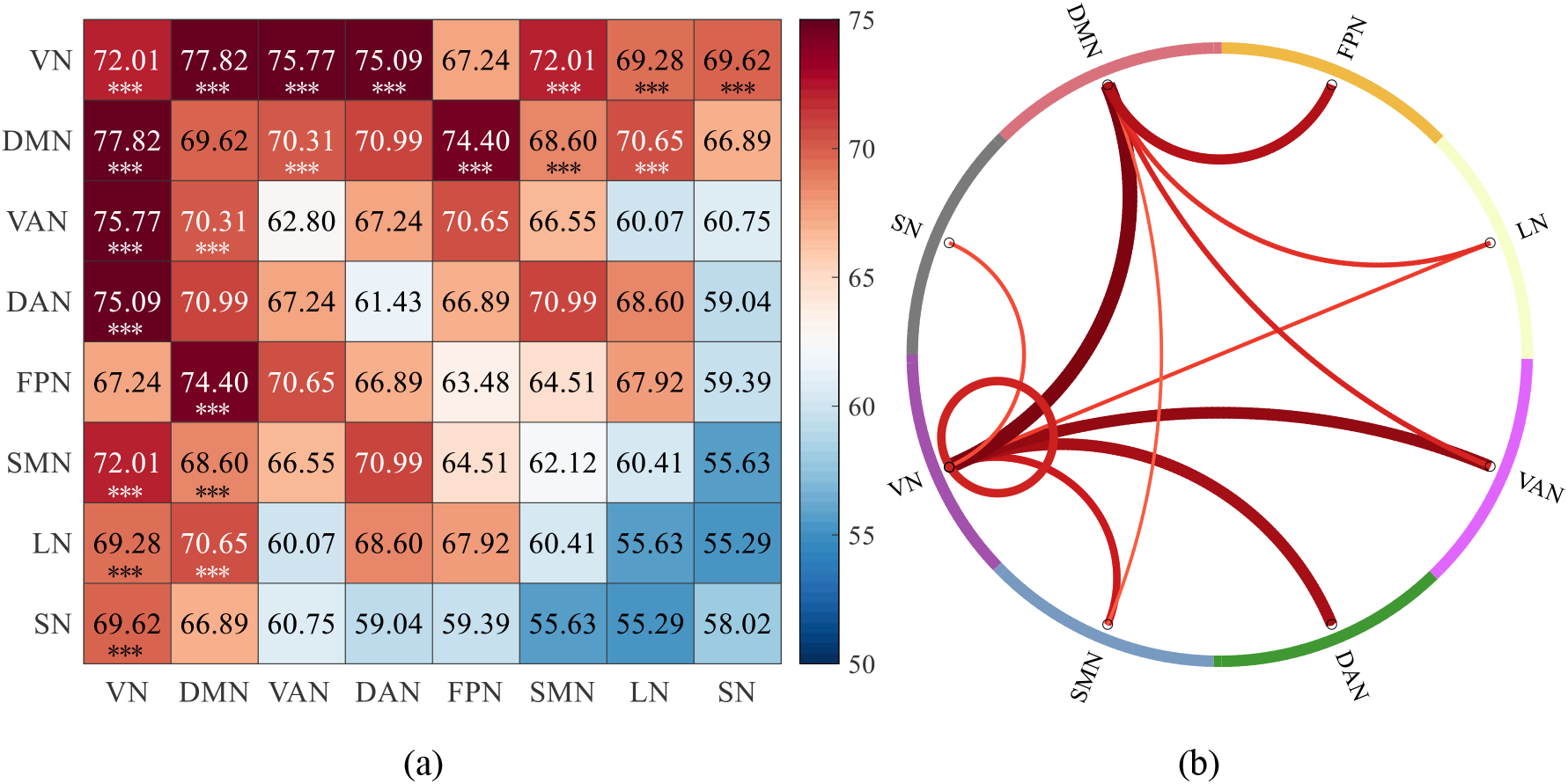
The network-based emotion prediction performance. (a) The prediction accuracy when each network is separately used for modeling. Here, *** indicates the networks with statistically significant predictive ability (p<0.0001, FDR corrected). (b) The VN and DMN related network-based functional connections with a statistically significant predictive ability (p<0.0001, FDR corrected). Thicker and redder connection lines indicate higher classification accuracy and vice versa.

### Subnetwork-based emotion prediction

The above network-based classification results suggest that the differences between happiness and sadness are primarily located in the functional connections within and between the VN and DMN. Next, we further investigate the details in the corresponding networks and analyze the predictive abilities of the involved subnetworks. Examination of functional networks in terms of subnetworks is expected to offer a deeper understanding of the predictive ability in the distinction of the two distinct basic emotions. According to the defined 18 large-scale subnetworks (Schaefer et al., 2018), the VN is divided into VN-a (central visual area) and VN-b (peripheral visual area); and DMN is divided into DMN-a, DMN-b and DMN-c. Besides, the VN and DMN related networks are also separated into a number of subnetworks. Here, VAN is divided into VAN-a and VAN-b; DAN is divided into DAN-a and DAN-b; and FPN is divided into FPN-a, FPN-b and FPN-c. Based on the subnetwork connections, we estimate the corresponding prediction performance on the distinction of emotions (Fig. 5 (a)). After the permutation test, the functional connections with significant predictive accuracies (p<0.0001, FDR corrected) are mainly distributed within DMN, between DMN and other subnetworks, and between VN-a and other subnetworks, as shown in Fig. 5 (b). All the obtained significance of the prediction results relative to the random level is reported in Appendix III of Supplementary Materials. After the statistical analysis, the significant predictive subnetworks with detailed performance are reported in Table 2. Here, the best prediction performance is achieved when the FC between DMN-a and FPN-c is utilized (*P_acc_*=74.40%, *P_f_* =75.08%, *P_pre_* =73.38%, *P_sen_* =76.87%, *P_spe_* =71.92%). Following, the other significant prediction performance is also observed when the FC patterns of VN-a - VN-b, VN-a - DMN-c, VN-a - DMN-a, VN-a - DAN-b, VN-a - VAN-b, VN-a - DAN-a, VN-a - LN-a, DMN-b - DAN-a, DMN-b - DMN-c, DMN-a - DMN-c, VN-a - FPN-a, DMN-a - VAN-a are used. All the above-reported classification accuracies are significantly greater than the chance level (p<0.0001, FDR corrected). It is found that the classification performance using the functional connections related to the central visual area is better than that using the functional connections related to the surrounding visual area. This indicates that the predictive functional connection patterns in the visual network mainly exist in the central visual area, instead of the surrounding visual area. For the functional connection between DMN and FPN, the most predictive patterns contributing to the emotion prediction are mainly distributed between DMN-a and FPN-c.

**Figure 5.**
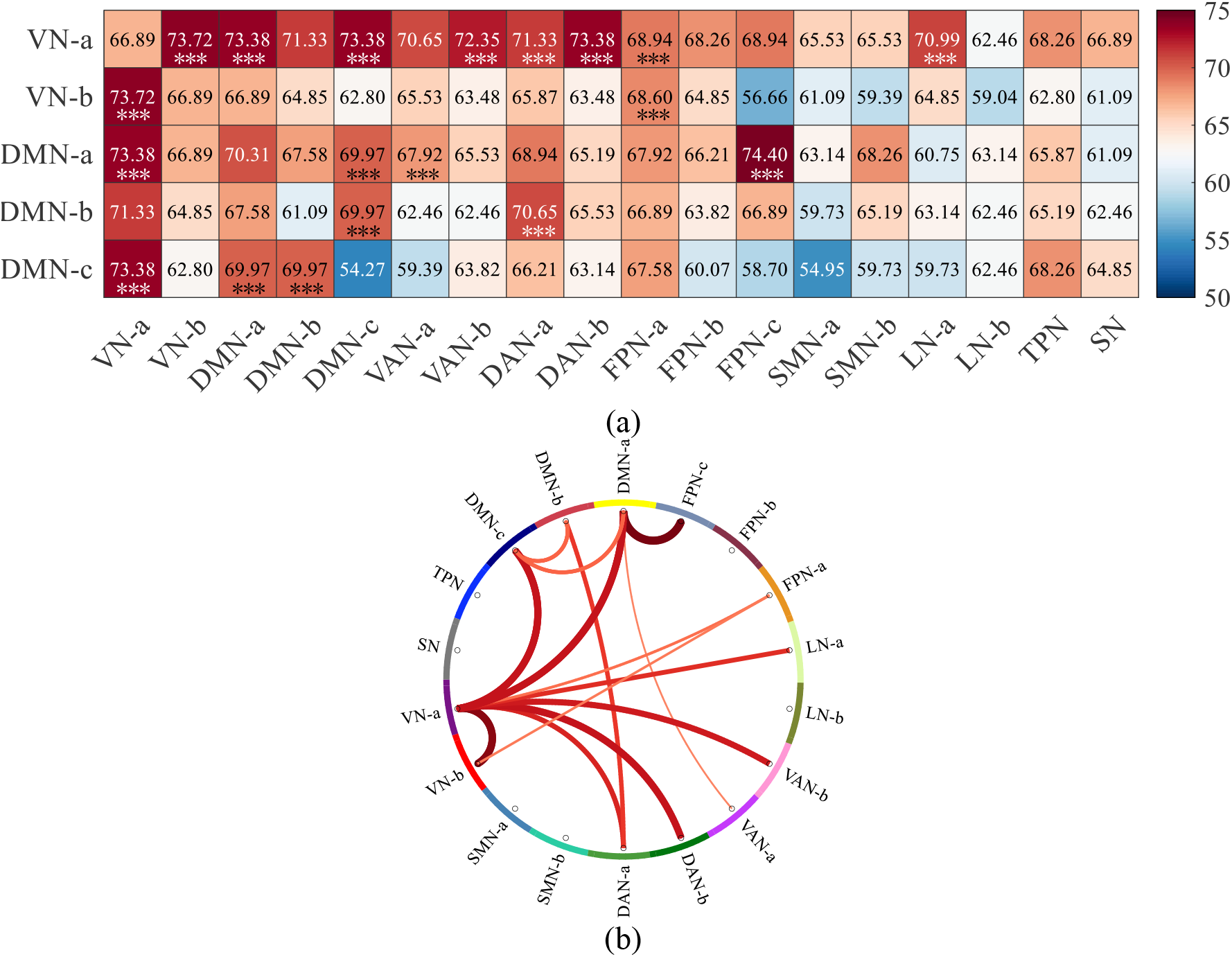
The classification accuracy of subnetwork-based emotion prediction. (a) The VN and DMN related subnetwork-based classification results. (b) The VN and DMN related subnetwork-based functional connections with a statistically significant predictive ability (p<0.0001, FDR corrected). Thicker and redder connection lines indicate higher classification accuracy and vice versa.

**Table 2.**
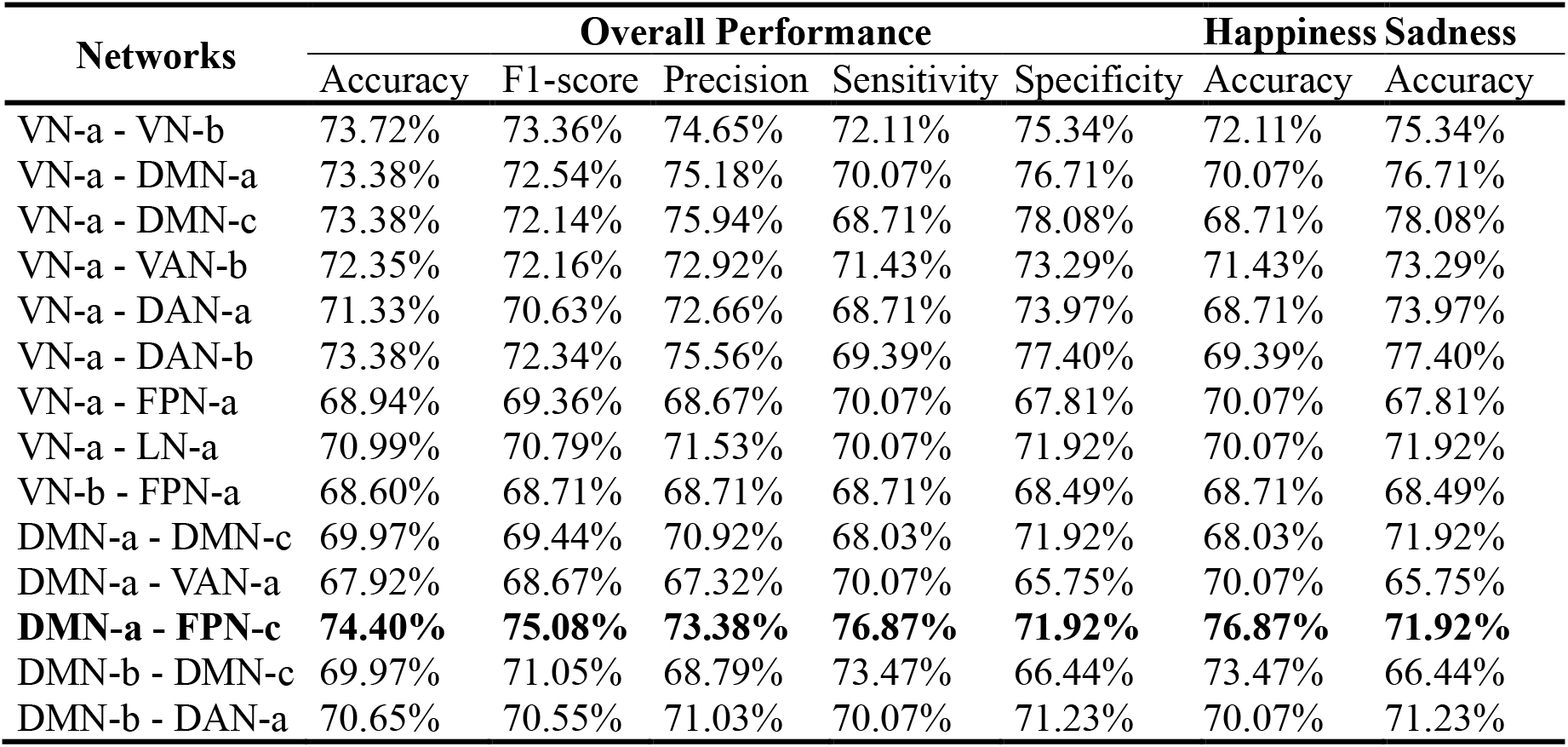
The predictive subnetworks with the detailed classification performance. All the reported classification results are statistically significant (p<0.0001, FDR corrected).

### Stimulation-stage-based emotion prediction

We further examined whether the FC profiles change over the period of sustained emotional experiences. To this end, we divide each episode into three stages: early stimulation (1 ~ 200 s), middle stimulation (201 ~ 400 s), and late stimulation (401 ~ 600 s), and perform the calculation of FCs at each period separately. For each stimulation stage, the emotion prediction models are separately built based on all VN and DMN related FCs, and the efficient emotion-evoking stage(s) are investigated in terms of emotion predictive ability. As shown in Table 3, the average prediction accuracies based on the entire stimulation period, early stimulation stage, middle stimulation stage, and late stimulation stage are 80.55% (*P_f_*=80.55%, *P_pre_*=80.82%, *P_sen_*=80.27%, *P_spe_*=80.82%), 78.50% (*P_f_* =77.89%, *P_pre_* =80.43%, *P_sen_* =75.51%, *P_spe_* =81.51%), 81.23% (*P_f_* =81.36%, *P_pre_* =81.08%, *P_sen_* =81.63%, *P_spe_* =80.82%), and 85.32% (*P_f_* =85.62%, *P_pre_* =84.21%, *P_sen_* =87.07%, *P_spe_* =83.56%), respectively. The prediction results between entire and late and between early and late are statistically different (p<0.05, FDR corrected). The corresponding confusion matrices are shown in Fig. 6. The results reveal that the classification performance of the entire stimulation period could be considered as an average performance of the early, middle, and late stimulation stages. Among the three stimulation stages, it is found that the prediction performance using the late stimulation period is superior to the other stimulation periods, which may additionally reflect that emotions evolve cumulatively over sustained exposure and long-term emotion stimulation could be beneficial to intense emotion elicitation.

**Table 3.**
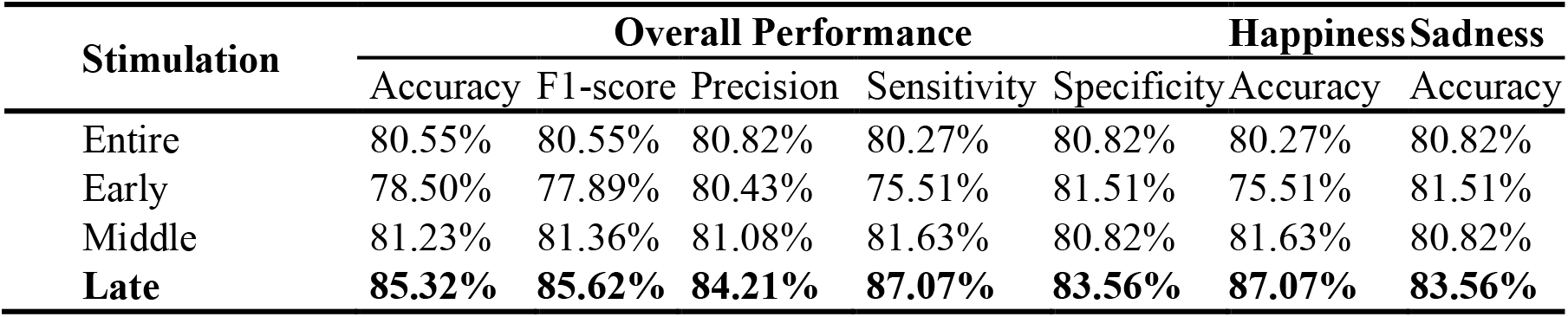
The predictive performance at different stimulation stages using all the VN and DMN related FCs with the detailed classification performance. All the results are statistically significant with p<0.0001 (FDR corrected).

**Figure 6.**
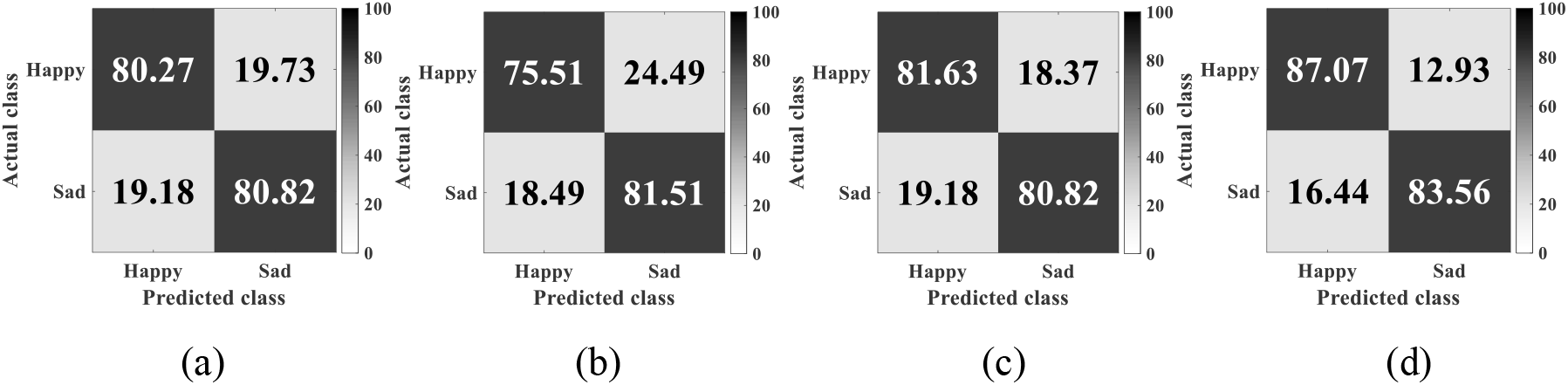
The obtained confusion matrices using all the VN and DMN related FCs at (a) the whole stimulation period, (b) the early stimulation stage, (c) the middle stimulation stage, and (d) the late stimulation stage.

Further, we estimate the predictive ability of FC patterns at each stimulation stage and examine whether there exists an FC pattern shifting of emotion-related contributions of the brain areas during the long-term emotional processing. All the obtained significance of the prediction results relative to the random level is reported in Appendix IV of Supplementary Materials. As shown in Fig. 7, it is found the predominantly distributed emotion-related FC patterns are slightly different at different stimulation stages. For the early stimulation stage, the emotion-related FC patterns are predominantly distributed in DMN-a - FPN-a (*P_acc_*=72.01%), VN-a - VN-b (*P_acc_*=71.33%), and VN-a - DAN-a (*P_acc_*=71.33%). For the middle stimulation stage, the emotion-related FC patterns are predominantly distributed in VN-a - SN (*P_acc_*=73.38%), VN-a - DMN-b (*P_acc_*=72.70%), VN-b - DAN-a (*P_acc_*=72.35%), DMN-a - DMN-b (*P_acc_*=72.01%), VN-a - DAN-a (*P_acc_*=71.67%), VN-a - VN-b (*P_acc_*=71.33%), VN-a - DAN-b (*P_acc_*=71.33%), VN-a - VAN-a (*P_acc_*=70.99%), and VN-a - LN-a (*P_acc_* =70.99%). For the late stimulation stage, the emotion-related FC patterns are predominantly distributed in VN-a and VN-b (*P_acc_*=72.01%), VN-a and VAN-b (*P_acc_*=72.01%), DMN-a and DMN-b (*P_acc_*=70.99%). On the other hand, all the predictive results using a single network are much lower than those using all the VN and DMN related FCs. The results demonstrate that emotion-related brain functions are dominated by distributed network systems, instead of a single network/area.

**Figure 7.**
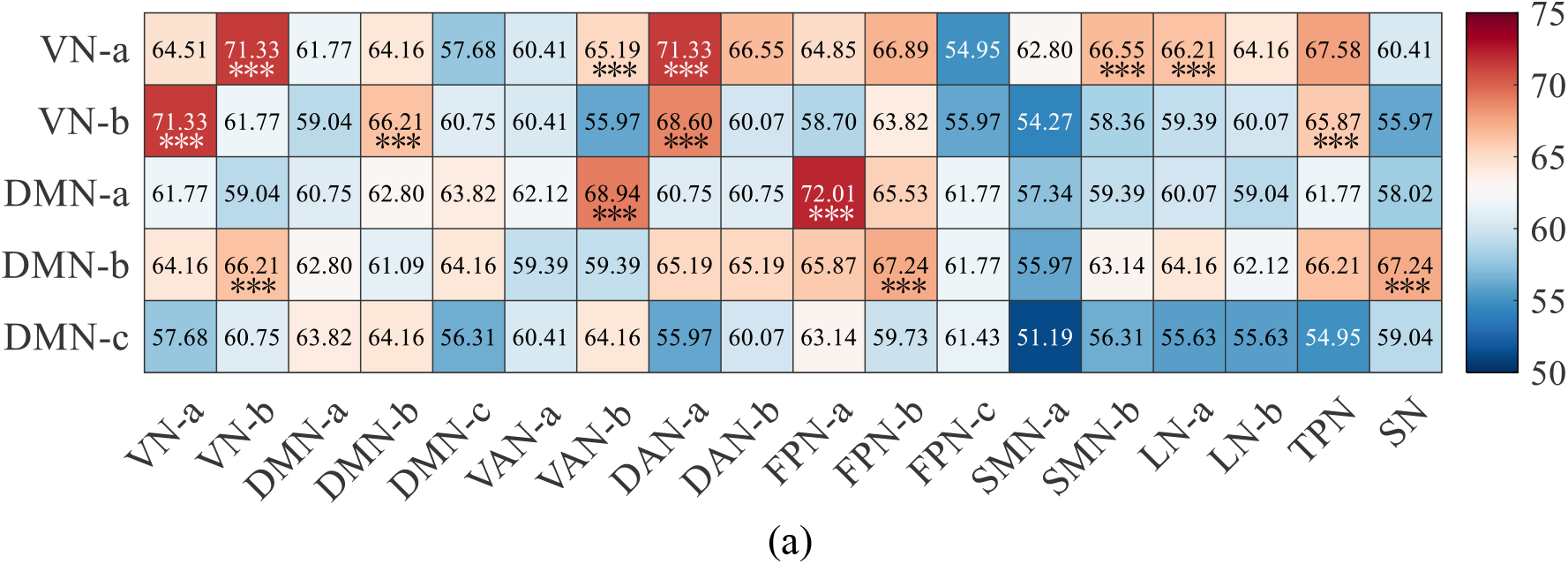
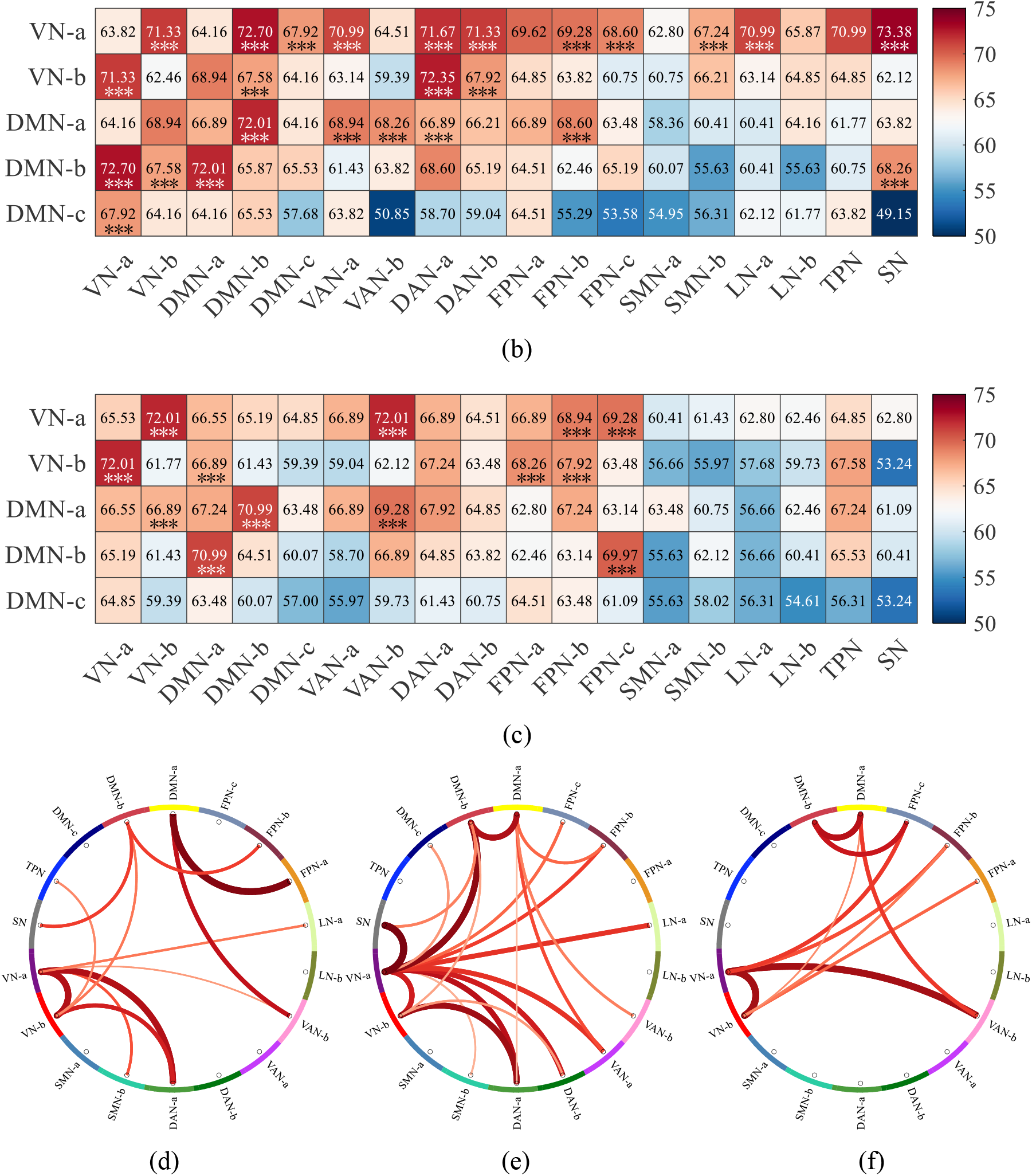
The obtained predictive abilities of FC patterns at different stimulation stages: (a) early, (b) middle, and (c) late. The corresponding subnetwork-based functional connections with statistically significant predictive ability (p<0.0001, FDR corrected) are shown in (d), (e), and (f). Thicker and redder connection lines indicate higher classification accuracy and vice versa.

## Discussion

The present study capitalized on sustained and ecologically valid induction of happiness and sadness via long movie clips and applied predictive modeling to determine whether these emotions are represented in distinct network level signatures. We therefore explored high resolution network level profiles and developed neuroimaging-based cross-subject cross-episode emotion prediction signatures via functional networks. The main objectives of the present study include: (1) applying the machine learning method to analyze fMRI FC information and providing a proof-of-concept emotion prediction model under a naturalistic experimental design employing sustained emotional induction, and (2) exploring distinct network patterns associated with two distinct basic emotions through a more natural experimental paradigm. We observed a significant difference in the whole-brain FC patterns when different emotions were evoked, where the results were fully evidenced by significantly higher emotion prediction performance than the random level (p<0.0001, FDR corrected). The results showed that the distinct basic emotions, happiness and sadness, have distinguishable and distributed neural representations on the whole-brain connectivity level, with the VN and DMN associated networks making the major contribution to the identification of the specific emotional states. Specific findings include (1) Happiness and sadness elicited by long-term movie clips have discrete neural representations, reflected in FC profiles. (2) The distinctive FC patterns of happiness and sadness are mainly distributed at VN and DMN associated networks. (3) Examining changes over the stimulation period (early, middle, and late presentation periods of the movie) reveal that emotional experience is an accumulative process such that the emotion-specific signatures become more distinct over the course of the sustained stimulation. (4) The estimated distinguishable ability of FC profiles on the sustained happiness and sadness are consistent across subjects and episodes. Together, these results underscore that interactions between brain regions contribute to emotional experiences under naturalistic conditions and that different emotions are represented in distinct network level profiles.

### Functional connectivity profiles associated with emotions

The present study shows that different patterns of whole-brain FC characterize specific emotional experiences. The emotion prediction results suggest that the whole-brain FC patterns can accurately distinguish between happy and sad emotions and that the corresponding emotion-specific changes are not restricted to a single region/network, but across multiple networks that primarily involve the VN and DMN. The VN and DMN may thus represent key network systems that encode specific emotional experiences and vary their interaction with other systems according to the external environment and the subjective emotional state. These findings converge with previous neuroimaging emotion studies which demonstrated the important roles of the VN and DMN networks during processes involving emotional experiences and emotion regulation (Jaworska et al., 2015; Nguyen et al., 2019; Phan et al., 2002; Satpute and Lindquist, 2019; Vytal and Hamann, 2010).

A previous fMRI study of emotion-related processing with visual stimuli revealed that the modulated brain regions in emotional processing critically relied on the stimulus type (Keightley et al., 2003). The superior classification performance between the visual network and other networks in the present study further supports the idea and suggests an important role in visual processing regions as early processing nodes for emotional information. Recent visual-related fMRI studies suggest that the role of visual processing areas extends beyond the simple perception of visual information (Cai et al., 2017; Guo et al., 2012; Katzner and Weigelt, 2013). A number of studies have shown that emotional content is also encoded and recovered in visual areas (ventral visual stream). For example, Mickley et al. (Mickley and Kensinger, 2008) found that, in the emotion encoding process, the involvement of visual areas made it possible for negatively related emotional memories to be recalled vividly. Kragel et al. (Kragel et al., 2019) observed that human visual cortical activity patterns enriched the encoding and decoding process of emotions. In addition, it has been found that different visual cortical areas are modulated by different categories of emotions (Thakral et al., 2022). Mourao-Miranda et al. (Mourao-Miranda et al., 2003) found that the pictures with negative emotions produced stronger activity in V1, compared to the pictures with positive emotions. In our subnetwork-based emotion prediction results, we observed that the emotion-related FC patterns in the visual areas are predominantly distributed in the central visual area (VN-a), instead of the peripheral visual area (VN-b). The central visual area includes the striate cortex (V1) and extrastriate cortex, while the peripheral visual area covers the extrastriate superior and inferior. One possible reason for the predominant distribution of the VN-a might be that emotions such as sadness involve stronger responses in sensory processing in the V1 cortex. One study has elucidated this phenomenon by suggesting that negative information elicits selective attentional priorities and attentional resources relative to positive information (Yiend, 2010).

The strong predictive abilities of DMN in the distinction of emotions align with previous studies. The DMN, including posterior cingulate cortex (PCC), precuneus, medial prefrontal cortex (MPFC), inferior parietal lobule (IPL), and bilateral temporal cortex regions, encompasses important and unique mental capacities (Raichle, 2015; Raichle et al., 2001; Satpute and Lindquist, 2019). DMN is found to be involved in internal attention, such as autobiographical memory (Buckner et al., 2008; Schacter and Addis, 2007), rumination (Hamilton et al., 2015; Whitfield-Gabrieli and Ford, 2012), social cognition (Spreng et al., 2009; Van Overwalle, 2009), social evaluation (Gusnard et al., 2001; Hamilton et al., 2015), and internal mentation (Andrews-Hanna et al., 2014). The neural substrates related to the internal sensory processing also contribute to emotional processing (Craig, 2009; Critchley et al., 2005; Pollatos et al., 2007). Previous studies also reported that the DMN plays a crucial role in the representation of individual emotions (Satpute and Lindquist, 2019), especially the involved ventral and anterior medial prefrontal cortices (vmPFC and amPFC). As the central areas in DMN, vmPFC, and amPFC are associated with emotion generation, integration, processing, and regulation (Gusnard et al., 2001; Raichle et al., 2001; Veer et al., 2011). Satpute and Lindquist also suggested that DMN may participate in emotion by supporting conceptual progress, which facilitates the ability to experience specific physiological sensations and contribute to the composition of emotion categories (Satpute and Lindquist, 2019). The present study confirms that DMN supports the integration of both visual and semantic information (Lim et al., 2013) and the involvement in emotion regulation and emotion-related decision-making (Rolls et al., 2022).

### Emotion-related coordinated function of distributed network systems

Using the whole-brain FC profiles for emotion prediction achieves a significantly better performance than using single network information. This suggests that emotion-specific FC patterns are not present in a single network or region, but require a distributed representation within and between multiple networks. In the investigation of the emotion-associated brain circuits, it was observed that emotion relies on large-scale functional network interaction (Pessoa, 2017). Studying the whole-brain FC using multivariate and machine learning analysis benefits the prediction of emotions (Pessoa, 2018). The powerful predictive ability of VN and DMN associated network systems supports that the neural representation of different emotional experiences is a distributed representation with VN and DMN as the core and multiple networks operating in concert. The involvement of large-scale brain interaction provides evidence that the connections between brain regions play a significant role in emotion.

These observations align with previous emotion studies, reporting that a number of fundamental emotional states depend on large-scale cortical and subcortical interactions, rather than engaging isolated networks/regions (Damasio and Carvalho, 2013; Kober et al., 2008; Lindquist et al., 2012; Nummenmaa et al., 2014; Saarimäki et al., 2016; Vytal and Hamann, 2010). Together with the present findings, this reflects that emotions are represented in distributed networks spanning multiple brain systems (Kragel and LaBar, 2015; Wager et al., 2015). It was found the predictive voxels for emotion decoding showed a similar tendency to distribute across the canonical networks in which VN and DMN networks accounted for about half of the total and the remaining DAN, VAN, LN, FPN, and SMN networks accounted for about the other half. Wager et al. (Wager et al., 2015) used a hierarchical Bayesian model to analyze the patterns of human brain activity under five emotion categories (fear, anger, disgust, sadness, and happiness), showing that emotion categories were not contained in any one region or system but were represented as synergistic across multiple brain networks cooperation. Saarimäki et al.’s work (Saarimäki et al., 2016) also verified the neural representations of discrete emotional experiences are distributed across multiple brain regions (Gao et al., 2020; Horikawa et al., 2020; Saarimäki et al., 2018). Different emotions are associated with activation changes across multiple functional systems, and the underlying spatial distribution configuration ultimately defines specific emotions at the psychological and behavioral levels (Saarimäki et al., 2018). Furthermore, the theory of constructed emotion suggests that all emotions consist of a shared set of basic functional systems that are not specific to emotion processing per se (Kober et al., 2008; Lindquist et al., 2012). We consider that the shared basic emotional systems consisting of discrete emotional experiences induced by the long-term naturalistic continuous emotion-evoking paradigm exhibit a similar engagement of networks at the whole-brain level, yet that VN and DMN may play an integrative role within these networks.

### Engagement of brain networks during sustained naturalistic emotional experience

Naturalistic stimulation, such as audio narratives or short movies, has been increasingly demonstrated to allow a more ecologically valid and comprehensive experimental assessment of brain processes than sparse experimental stimuli (e.g. emotional words) with high reproducibility (Matusz et al., 2019; Zhang et al., 2021). Compared with the traditional experimental paradigms of blocked design or event-related design, naturalistic stimulation shows some compelling advantages (Bottenhorn et al., 2018; Meer et al., 2020; Simony and Chang, 2020) including the engagement of complex and interacting brain states while closely simulating brain processes in real-life (DuPre et al., 2020; Jääskeläinen et al., 2021). Emerging evidence and conceptual work suggest that ecologically valid scenarios offer some benefits over traditional parametric task designs, including a test of experimental brain models under ecologically valid conditions (Puckett et al., 2020; Sonkusare et al., 2019; Vanderwal et al., 2022). With respect to emotion-related studies, the naturalistic paradigms enable a more ecologically valid approach to the dynamic neurophysiological processes that underly the different emotional states in everyday life (Lettieri et al., 2019).

However, most of the previous emotion-related studies are conducted using naturalistic stimuli with a short duration, such as pictures (Bush et al., 2018), music (Putkinen et al., 2021), short movie clips (Wang et al., 2017a), and movie trailers (Chan et al., 2020), to elicit subjects’ specific emotional experience in a highly controlled laboratory environment. In real life, several emotional states evolve over longer time periods and emotions such as vivid sadness may require a high level of contextualization to fully evolve. The emotion elicitation with short stimuli may fail to effectively evoke the specific emotional experience, which would further affect the following brain analysis and lead to poor emotion estimation results. For example, in Saarimäki et al.’s work (Saarimäki et al., 2022), the mean classification accuracy on sadness was the lowest (18%), which was close to the chance level (16.67%). A similar situation occurred in another fMRI-based emotion recognition work (Saarimäki et al., 2016), where the emotion prediction result on sadness was still the lowest. One possible reason for leading the classification performance on sadness being much worse than the other emotions could be that the stimulus ((Saarimäki et al., 2022): 1-minute narrative; (Saarimäki et al., 2016): 10-second movie clips) is too short to elicit a strong and deep sad emotion and further fail to bring a significant change in the brain activities. The previous results suggest that effective emotion elicitation needs a longer time, especially for sad emotional experience.

Betzel et al. (Betzel et al., 2020) found the temporal fluctuation in network integration and segregation during movie-watching, sharing consistent patterns across individuals. In the present study, we divide the whole stimulation period into early, middle, and late stages to explore whether the network level representations change over time during sustained naturalistic emotion processing. We found that the corresponding prediction performance at the late stimulation stage (84.71%) was considerably better as compared to the early (78.17%) and middle (80.49%) stimulation stages, while the decoding performance for both, happy and sad emotional experience was generally high. As evidenced by the stimulation-stage-based emotion prediction results, the emotional experience is sustainable accumulation, reflected in the corresponding brain activities with evidence of the growth predictive ability. This resonates with previous findings suggesting that the time of emotional exposure may affect the emotional experiences and the neural expressions (Résibois et al., 2017; Waugh and Kuppens, 2021; Waugh et al., 2012). For instance, emotional stimuli tend to be better remembered in long-lasting contextual memory and those long-term emotional stimuli can enhance the effectiveness of emotional elicitation by enhancing contextual memory (Dolcos et al., 2013).

### Predictive ability is consistent across subjects and movie clips

Through a comprehensive network level description of whole-brain FC patterns may allow a powerful predictive determination of specific emotional states such as happiness and sadness. Further, we examine the consistency of the predictive ability on different subjects and movie clips, respectively.

To explore the individual differences in prediction performance, we separately examine each subject’s decoding performance to help clarify the consistency of emotion-related contributions of brain networks. Strong predictive abilities of VN and DMN associated networks are consistently observed across subjects. However, there still exists a variance in the prediction performance among different individuals, which may be caused by the different sensitivities of emotional experience and emotional perception across subjects. We discover that the emotion prediction on subject 32 achieves a good performance, where the vast majority of the classification accuracies within or between networks are greater than 70%. In contrast, the prediction performance on subject 24 is close to the chance level. These observations underscore individual differences and may reflect different levels of emotional engagement between the subjects or individual variations in the “typical” brain organization. The difference in emotion perception may be caused by the nature of cognitive processes that have different emotion sensitivities to the stimulation, rather than the properties of the stimuli themselves (Dolcos et al., 2013; Petro et al., 2018).

In addition, to verify the emotion stimulation effect of the selected movie clips, we also investigate the cross-subject prediction performance of each movie clip using different intra- and inter-networks. It is found that consistent good prediction performance can be observed across movie clips when VN and DMN associated networks are adopted. The highest predictive ability is based on the connections between VN and DMN, where the prediction accuracies of all movie clips are larger than 70%. These results show that the predictive ability of FC patterns is highly consistent across different movie clips.

## Conclusion

The present study determined the predictive properties of whole-brain FC patterns to determine distinct neurofunctional representations of two distinct basic emotions during naturalistic movie watching. This approach allowed us to explore a higher-order neural process under ecologically valid experimental conditions of sustained emotional experiences. Our work provides preliminary evidence that the VN and DMN associated networks exhibit a strong contribution to the prediction and thus may represent integrative networks that orchestrate the whole-brain network level expressions of specific emotional states. The results further emphasize the role of network level changes as the basis of different sustained emotional experiences under ecologically valid conditions. Our findings reveal that the naturalistic long-term movie-watching paradigm evoked emotions manifest as distributed representations of multiple networks operating in concert with the VN and DMN as the core. This paper provides compelling evidence and unique insights into the emotion-related FC patterns that supports emotion perceiving and processing and reveals the importance of the VN and DMN coordination to emotions.

## Acknowledgments

This work was supported in part by the National Natural Science Foundation of China under Grants 62276169, 61906122, 32250610208, and 82271583, in part by Shenzhen-Hong Kong Institute of Brain Science-Shenzhen Fundamental Research Institutions (2022SHIBS0003), in part by the Tencent “Rhinoceros Birds”-Scientific Research Foundation for Young Teachers of Shenzhen University, and in part by the High-Level University Construction under Grant 000002110133.

