## Supplementary Material for "Whole-Brain fMRI Functional Connectivity Signatures Predict Sustained Emotional Experience in Naturalistic Contexts"

### Supplementary Materials

#### Appendix I: Stimuli Information

In this work, a total of 12 movie clip candidates were selected and edited for emotion evoking. The edited source information is listed in Table S-1. To avoid selection bias, we invited volunteers to select the proper stimuli for the emotion experiment. Only the video clips with consistent emotion feedback from different volunteers were included. Each video clip candidate was assessed at least by 20 volunteers, and there was no overlap between the volunteers (selecting proper stimuli) and the subjects (participating in the fMRI experiment). During the fMRI experiment, we also collected the subjects' feedback after each episode based on a 5-point rating scale. The instruction words were: please rate your emotion state when you were watching this video as very happy [10] / happy [5] / neutral [0] / sad [-5] / very sad [-10]. The collected emotion feedback of each movie clip candidate from the subjects is presented in Table S-2, which verify the selected movie clips could successfully evoke the intended emotions effectively and reliably (the feedbacks are larger than 5 for happy stimuli and are smaller than -5 for sad stimuli).

Table S-1. The edited source information of the selected stimuli. The numbers 1~6 are the edited movie clips for happiness, and 7~12 are the edited movie clips for sadness.

| **No.** | **Movie Name** | **Start-End** | **No.** | **Movie Name** | **Start-End** |
|---|---|---|---|---|---|
| 1 | Mr. Popper's Penguins | 0:17:59-0:27:59 | 2 | Ted | 0:01:06-0:11:06 |
| 3 | The Onion Movie | 0:00:44-0:10:44 | 4 | Liar Liar | 0:38:16-0:48:16 |
| 5 | A Thousand Words | 0:22:17-0:32:17 | 6 | Absolutely Anything | 0:20:05-0:30:05 |
| 7 | Miracle In Cell No.7 | 1:20:49-1:30:49 | 8 | Prayers For Bobby | 0:39:07-0:49:07 |
| 9 | The Classic | 1:50:42-2:00:42 | 10 | Grave Of The Fireflies | 1:10:09-1:20:09 |
| 11 | Only The Brave | 1:51:43-2:01:43 | 12 | The Last Train | 1:24:33-1:34:33 |

Table S-2. The collected subjects' feedback on emotion properties. The numbers 1~6 are the edited movie clips for happiness, and 7~12 are the edited movie clips for sadness.

| **No.** | **Feedback** | **No.** | **Feedback** |
|---|---|---|---|
| **1** | 5.2±3.3 | **2** | 5.8±2.7 |
| **3** | 5.2±3.4 | **4** | 6.1±2.6 |
| **5** | 5.3±2.7 | **6** | 3.1±2.9 |
| **Average feedback on all the selected happy movie clips:** | | | **5.2±3.0** |
| **7** | -5.2±2.6 | **8** | -6.1±3.3 |
| **9** | -6.0±3.0 | **10** | -2.9±2.5 |
| **11** | -4.5±3.1 | **12** | -5.2±3.3 |
| **Average feedback on all the selected sad movie clips:** | | | **-5.1±3.2** |

### Appendix II: Network-Based Emotion Prediction

The significant network-based emotion prediction results after false discovery rate (FDR) correction is shown in Table S-3. Here, we consider the prediction results of both happiness and sadness as greater than the random prediction results as significant (p<0.0001, FDR corrected).

Table S-3. The obtained p-values of the network-based emotion prediction after FDR correction. The highlights are the intra- and inter-networks with a significant prediction of both happiness and sadness (p<0.0001, FDR corrected).

| Networks | Overall | Happiness | Sadness | Networks | Overall | Happiness | Sadness |
|---|---|---|---|---|---|---|---|
| VN | <0.0001 | <0.0001 | <0.0001 | DAN-FPN | <0.0001 | 0.0033 | <0.0001 |
| VN-SMN | <0.0001 | <0.0001 | <0.0001 | DAN-DMN | <0.0001 | 0.0033 | <0.0001 |
| VN-DAN | <0.0001 | <0.0001 | <0.0001 | DAN-SN | <0.0001 | 0.0664 | 0.0418 |
| VN-VAN | <0.0001 | <0.0001 | <0.0001 | VAN | <0.0001 | <0.0001 | 0.0286 |
| VN-LN | <0.0001 | <0.0001 | <0.0001 | VAN-LN | <0.0001 | 0.0019 | 0.0921 |
| VN-FPN | <0.0001 | 0.0047 | <0.0001 | VAN-FPN | <0.0001 | <0.0001 | 0.0015 |
| VN-DMN | <0.0001 | <0.0001 | <0.0001 | VAN-DMN | <0.0001 | <0.0001 | <0.0001 |
| VN-SN | <0.0001 | <0.0001 | <0.0001 | VAN-SN | <0.0001 | 0.0472 | 0.0093 |
| SMN | <0.0001 | 0.0552 | 0.0043 | LN | 0.0310 | 0.0557 | 0.4250 |
| SMN-DAN | <0.0001 | <0.0001 | 0.0015 | LN-FPN | <0.0001 | <0.0001 | 0.0015 |
| SMN-VAN | <0.0001 | 0.0130 | <0.0001 | LN-DMN | <0.0001 | <0.0001 | <0.0001 |
| SMN-LN | <0.0001 | 0.0402 | 0.0360 | LN-SN | 0.0169 | 0.1186 | 0.1687 |
| SMN-FPN | <0.0001 | 0.0120 | <0.0001 | FPN | <0.0001 | 0.0709 | 0.0015 |
| SMN-DMN | <0.0001 | <0.0001 | <0.0001 | FPN-DMN | <0.0001 | <0.0001 | <0.0001 |
| SMN-SN | 0.0175 | 0.2705 | 0.0484 | FPN-SN | <0.0001 | 0.0411 | 0.0687 |
| DAN | <0.0001 | 0.0411 | 0.0055 | DMN | <0.0001 | <0.0001 | 0.0015 |
| DAN-VAN | <0.0001 | 0.0033 | <0.0001 | DMN-SN | <0.0001 | <0.0001 | 0.0015 |
| DAN-LN | <0.0001 | 0.0019 | <0.0001 | SN | 0.0011 | 0.3450 | 0.0141 |

#### Appendix III: Subnetwork-Based Emotion Prediction

The significant subnetwork-based emotion prediction results after false discovery rate (FDR) correction is shown in Table S-4. Here, we consider the prediction results of both happiness and sadness as greater than the random prediction results as significant (p<0.0001, FDR corrected).

Table S-4. The obtained p-values of the subnetwork-based emotion prediction after FDR correction. The highlights are the intra- and inter-networks with a significant prediction of both happiness and sadness (p<0.0001, FDR corrected).

| Networks | Overall | Happiness | Sadness | Networks | Overall | Happiness | Sadness |
|---|---|---|---|---|---|---|---|
| VN-a | <0.0001 | 0.0110 | <0.0001 | DMN-a - VAN-b | <0.0001 | 0.0023 | 0.0106 |
| VN-a - VN-b | <0.0001 | <0.0001 | <0.0001 | DMN-a - LN-b | <0.0001 | 0.0070 | 0.0377 |
| VN-a - SMN-a | <0.0001 | 0.0098 | <0.0001 | DMN-a - LN-a | <0.0001 | 0.0098 | 0.1091 |
| VN-a - SMN-b | <0.0001 | 0.0124 | 0.0022 | DMN-a - FPN-a | <0.0001 | 0.0040 | 0.0022 |
| VN-a - DAN-a | <0.0001 | <0.0001 | <0.0001 | DMN-a - FPN-b | <0.0001 | 0.0057 | 0.0050 |
| VN-a - DAN-b | <0.0001 | <0.0001 | <0.0001 | DMN-a - FPN-c | <0.0001 | <0.0001 | <0.0001 |
| VN-a - VAN-a | <0.0001 | 0.0023 | <0.0001 | DMN-a | <0.0001 | <0.0001 | 0.0038 |
| VN-a - VAN-b | <0.0001 | <0.0001 | <0.0001 | DMN-a - DMN-b | <0.0001 | <0.0001 | 0.0063 |
| VN-a - LN-b | <0.0001 | 0.0246 | 0.0038 | DMN-a - DMN-c | <0.0001 | <0.0001 | <0.0001 |
| VN-a - LN-a | <0.0001 | <0.0001 | <0.0001 | DMN-a - TPN | <0.0001 | 0.0040 | <0.0001 |
| VN-a - FPN-a | <0.0001 | <0.0001 | <0.0001 | DMN-a - SN | <0.0001 | 0.0246 | 0.0190 |
| VN-a - FPN-b | <0.0001 | 0.0023 | <0.0001 | DMN-b - SMN-a | <0.0001 | 0.0578 | 0.0147 |
| VN-a - FPN-c | <0.0001 | 0.0040 | <0.0001 | DMN-b - SMN-b | <0.0001 | 0.0023 | 0.0050 |
| VN-a - DMN-a | <0.0001 | <0.0001 | <0.0001 | DMN-b - DAN-a | <0.0001 | <0.0001 | <0.0001 |
| VN-a - DMN-b | <0.0001 | 0.0023 | <0.0001 | DMN-b - DAN-b | <0.0001 | 0.0124 | <0.0001 |
| VN-a - DMN-c | <0.0001 | <0.0001 | <0.0001 | DMN-b - VAN-a | <0.0001 | 0.0552 | 0.0050 |
| VN-a - TPN | <0.0001 | 0.0110 | <0.0001 | DMN-b - VAN-b | <0.0001 | 0.0116 | 0.0462 |
| VN-a - SN | <0.0001 | 0.0023 | 0.0022 | DMN-b - LN-b | <0.0001 | 0.0241 | 0.0129 |
| VN-b | <0.0001 | 0.0070 | <0.0001 | DMN-b - LN-a | <0.0001 | 0.0136 | 0.0147 |
| VN-b - SMN-a | <0.0001 | 0.0341 | 0.0200 | DMN-b - FPN-a | <0.0001 | 0.0023 | <0.0001 |
| VN-b - SMN-b | <0.0001 | 0.0570 | 0.0348 | DMN-b - FPN-b | <0.0001 | <0.0001 | 0.0715 |
| VN-b - DAN-a | <0.0001 | 0.0070 | 0.0022 | DMN-b - FPN-c | <0.0001 | 0.0023 | 0.0022 |
| VN-b - DAN-b | <0.0001 | 0.0219 | 0.0050 | DMN-b | <0.0001 | 0.0147 | 0.0526 |
| VN-b - VAN-a | <0.0001 | 0.0085 | <0.0001 | DMN-b - DMN-c | <0.0001 | <0.0001 | <0.0001 |
| VN-b - VAN-b | <0.0001 | 0.0184 | <0.0001 | DMN-b - TPN | <0.0001 | 0.0070 | 0.0138 |
| VN-b - LN-b | <0.0001 | 0.1116 | 0.0222 | DMN-b - SN | <0.0001 | 0.0267 | 0.0022 |
| VN-b - LN-a | <0.0001 | <0.0001 | 0.0168 | DMN-c - SMN-a | 0.0273 | 0.2036 | 0.1313 |
| VN-b - FPN-a | <0.0001 | <0.0001 | <0.0001 | DMN-c - SMN-b | <0.0001 | 0.0757 | 0.0138 |
| VN-b - FPN-b | <0.0001 | 0.0040 | 0.0022 | DMN-c - DAN-a | <0.0001 | <0.0001 | 0.0050 |
| VN-b - FPN-c | 0.0041 | 0.1364 | 0.1053 | DMN-c - DAN-b | <0.0001 | 0.0124 | 0.0119 |
| VN-b - DMN-a | <0.0001 | <0.0001 | 0.0063 | DMN-c - VAN-a | <0.0001 | 0.0377 | 0.0467 |
| VN-b - DMN-b | <0.0001 | 0.0116 | 0.0022 | DMN-c - VAN-b | <0.0001 | 0.0023 | 0.0230 |
| VN-b - DMN-c | <0.0001 | <0.0001 | 0.0616 | DMN-c - LN-b | <0.0001 | 0.0116 | 0.0129 |
| VN-b - TPN | <0.0001 | 0.0322 | 0.0063 | DMN-c - LN-a | <0.0001 | 0.0040 | 0.2197 |
| VN-b - SN | <0.0001 | 0.1099 | 0.0038 | DMN-c - FPN-a | <0.0001 | <0.0001 | 0.0022 |
| DMN-a - SMN-a | <0.0001 | 0.0116 | 0.0157 | DMN-c - FPN-b | <0.0001 | 0.0348 | 0.0318 |
| DMN-a - SMN-b | <0.0001 | <0.0001 | 0.0038 | DMN-c - FPN-c | 0.0010 | 0.1164 | 0.0239 |
| DMN-a - DAN-a | <0.0001 | 0.0023 | <0.0001 | DMN-c | 0.0460 | 0.2380 | 0.2260 |
| DMN-a - DAN-b | <0.0001 | 0.0057 | 0.0050 | DMN-c - TPN | <0.0001 | <0.0001 | 0.0038 |
| DMN-a - VAN-a | <0.0001 | <0.0001 | <0.0001 | DMN-c - SN | <0.0001 | 0.0023 | 0.0106 |

### Appendix IV: Stimulation-Stage-Based Emotion Prediction

The significant emotion prediction results after false discovery rate (FDR) correction at early, middle, and later stimulation stages are shown in Table S-5, Table S-6, and Table S-7, respectively. Here, we consider the prediction results of both happiness and sadness as greater than the random prediction results as significant ($p<0.0001$, FDR corrected).

Table S-5. The obtained p-values of the emotion prediction results after FDR correction at the early stimulation stage. The highlights are the intra- and inter-networks with a significant prediction of both happiness and sadness ($p<0.0001$, FDR corrected).

| Networks | Overall | Happiness | Sadness |
|---|---|---|---|
| VN-a | <0.0001 | 0.0105 | 0.0103 |
| VN-a - VN-b | <0.0001 | <0.0001 | <0.0001 |
| VN-a - SMN-a | <0.0001 | 0.0303 | 0.0030 |
| VN-a - SMN-b | <0.0001 | <0.0001 | <0.0001 |
| VN-a - DAN-a | <0.0001 | <0.0001 | <0.0001 |
| VN-a - DAN-b | <0.0001 | 0.0035 | <0.0001 |
| VN-a - VAN-a | <0.0001 | 0.0094 | 0.0988 |
| VN-a - VAN-b | <0.0001 | <0.0001 | <0.0001 |
| VN-a - LN-b | <0.0001 | 0.0257 | <0.0001 |
| VN-a - LN-a | <0.0001 | <0.0001 | <0.0001 |
| VN-a - FPN-a | <0.0001 | 0.0035 | 0.0055 |
| VN-a - FPN-b | <0.0001 | 0.0055 | <0.0001 |
| VN-a - FPN-c | 0.0267 | 0.3109 | 0.0758 |
| VN-a - DMN-a | <0.0001 | 0.0117 | 0.0280 |
| VN-a - DMN-b | <0.0001 | 0.0153 | <0.0001 |
| VN-a - DMN-c | 0.0024 | 0.1144 | 0.0450 |
| VN-a - TPN | <0.0001 | 0.0055 | <0.0001 |
| VN-a - SN | 0.0012 | 0.0153 | 0.0324 |
| VN-b | <0.0001 | 0.0130 | 0.0267 |
| VN-b - SMN-a | 0.0567 | 0.1952 | 0.1884 |
| VN-b - SMN-b | <0.0001 | 0.0459 | 0.0622 |
| VN-b - DAN-a | <0.0001 | <0.0001 | <0.0001 |
| VN-b - DAN-b | <0.0001 | 0.0105 | 0.0473 |
| VN-b - VAN-a | <0.0001 | 0.0055 | 0.1046 |
| VN-b - VAN-b | 0.0156 | 0.1425 | 0.1182 |
| VN-b - LN-b | <0.0001 | 0.0035 | 0.0955 |
| VN-b - LN-a | <0.0001 | 0.0105 | 0.1337 |
| VN-b - FPN-a | <0.0001 | 0.0167 | 0.1206 |
| VN-b - FPN-b | <0.0001 | <0.0001 | 0.0200 |
| VN-b - FPN-c | 0.0093 | 0.2255 | 0.0841 |
| VN-b - DMN-a | <0.0001 | 0.0204 | 0.0727 |
| VN-b - DMN-b | <0.0001 | <0.0001 | <0.0001 |
| VN-b - DMN-c | <0.0001 | 0.0204 | 0.0267 |
| VN-b - TPN | <0.0001 | <0.0001 | <0.0001 |
| VN-b - SN | 0.0200 | 0.0645 | 0.2203 |
| DMN-a - SMN-a | 0.0059 | 0.0962 | 0.0988 |
| DMN-a - SMN-b | <0.0001 | 0.0281 | 0.0549 |
| DMN-a - DAN-a | <0.0001 | 0.0077 | 0.1194 |
| DMN-a - DAN-b | <0.0001 | 0.0055 | 0.0757 |
| DMN-a - VAN-a | <0.0001 | 0.0105 | 0.0324 |
| DMN-a - VAN-b | <0.0001 | <0.0001 | <0.0001 |
| DMN-a - LN-b | 0.0012 | 0.0706 | 0.0622 |
| DMN-a - LN-a | <0.0001 | 0.0281 | 0.0409 |
| DMN-a - FPN-a | <0.0001 | <0.0001 | <0.0001 |
| DMN-a - FPN-b | <0.0001 | 0.0196 | <0.0001 |
| DMN-a - FPN-c | <0.0001 | 0.0094 | 0.0409 |
| DMN-a | <0.0001 | 0.0490 | 0.0165 |
| DMN-a - DMN-b | <0.0001 | 0.0055 | 0.0409 |
| DMN-a - DMN-c | <0.0001 | <0.0001 | 0.0267 |
| DMN-a - TPN | <0.0001 | 0.0035 | 0.0372 |
| DMN-a - SN | 0.0012 | 0.0267 | 0.0841 |
| DMN-b - SMN-a | 0.0184 | 0.2410 | 0.0585 |
| DMN-b - SMN-b | <0.0001 | 0.0591 | <0.0001 |
| DMN-b - DAN-a | <0.0001 | 0.0035 | 0.0030 |
| DMN-b - DAN-b | <0.0001 | 0.0117 | <0.0001 |
| DMN-b - VAN-a | 0.0012 | 0.0276 | 0.0892 |
| DMN-b - VAN-b | <0.0001 | 0.0153 | 0.1337 |
| DMN-b - LN-b | <0.0001 | <0.0001 | 0.0892 |
| DMN-b - LN-a | <0.0001 | 0.0035 | 0.0165 |
| DMN-b - FPN-a | <0.0001 | 0.0035 | 0.0030 |
| DMN-b - FPN-b | <0.0001 | <0.0001 | <0.0001 |
| DMN-b - FPN-c | <0.0001 | 0.0153 | 0.0165 |
| DMN-b | <0.0001 | 0.0215 | 0.0409 |
| DMN-b - DMN-c | <0.0001 | 0.0055 | 0.0030 |
| DMN-b - TPN | <0.0001 | <0.0001 | 0.0030 |
| DMN-b - SN | <0.0001 | <0.0001 | <0.0001 |
| DMN-c - SMN-a | 0.3170 | 0.3780 | 0.3990 |
| DMN-c - SMN-b | 0.0164 | 0.0594 | 0.1708 |
| DMN-c - DAN-a | 0.0146 | 0.1063 | 0.1845 |
| DMN-c - DAN-b | <0.0001 | 0.0117 | 0.0544 |
| DMN-c - VAN-a | <0.0001 | 0.1144 | <0.0001 |
| DMN-c - VAN-b | <0.0001 | 0.0094 | 0.0080 |
| DMN-c - LN-b | 0.0126 | 0.1827 | 0.1055 |
| DMN-c - LN-a | 0.0200 | 0.0380 | 0.3484 |
| DMN-c - FPN-a | <0.0001 | 0.0077 | 0.0030 |
| DMN-c - FPN-b | <0.0001 | 0.0525 | 0.0183 |
| DMN-c - FPN-c | <0.0001 | 0.0413 | 0.0055 |
| DMN-c | 0.0059 | 0.0525 | 0.2215 |
| DMN-c - TPN | 0.0267 | 0.2084 | 0.1206 |
| DMN-c - SN | 0.0012 | 0.0130 | 0.1194 |

Table S-6. The obtained p-values of the emotion prediction results after FDR correction at the middle stimulation stage. The highlights are the intra- and inter-networks with a significant prediction of both happiness and sadness ($p<0.0001$, FDR corrected).

| **Networks** | **Overall** | **Happiness** | **Sadness** | **Networks** | **Overall** | **Happiness** | **Sadness** |
|---|---|---|---|---|---|---|---|
| VN-a | <0.0001 | 0.0163 | 0.0019 | DMN-a - VAN-b | <0.0001 | <0.0001 | <0.0001 |
| VN-a - VN-b | <0.0001 | <0.0001 | <0.0001 | DMN-a - LN-b | <0.0001 | 0.0034 | 0.0123 |
| VN-a - SMN-a | <0.0001 | 0.0046 | 0.0117 | DMN-a - LN-a | <0.0001 | 0.0054 | 0.0730 |
| VN-a - SMN-b | <0.0001 | <0.0001 | <0.0001 | DMN-a - FPN-a | <0.0001 | <0.0001 | 0.0019 |
| VN-a - DAN-a | <0.0001 | <0.0001 | <0.0001 | DMN-a - FPN-b | <0.0001 | <0.0001 | <0.0001 |
| VN-a - DAN-b | <0.0001 | <0.0001 | <0.0001 | DMN-a - FPN-c | <0.0001 | <0.0001 | 0.0197 |
| VN-a - VAN-a | <0.0001 | <0.0001 | <0.0001 | DMN-a | <0.0001 | <0.0001 | 0.0145 |
| VN-a - VAN-b | <0.0001 | 0.0034 | <0.0001 | DMN-a - DMN-b | <0.0001 | <0.0001 | <0.0001 |
| VN-a - LN-b | <0.0001 | 0.0021 | <0.0001 | DMN-a - DMN-c | <0.0001 | 0.0021 | 0.0154 |
| VN-a - LN-a | <0.0001 | <0.0001 | <0.0001 | DMN-a - TPN | <0.0001 | 0.0452 | <0.0001 |
| VN-a - FPN-a | <0.0001 | 0.0021 | <0.0001 | DMN-a - SN | <0.0001 | 0.0034 | 0.0145 |
| VN-a - FPN-b | <0.0001 | <0.0001 | <0.0001 | DMN-b - SMN-a | <0.0001 | 0.0447 | 0.0187 |
| VN-a - FPN-c | <0.0001 | <0.0001 | <0.0001 | DMN-b - SMN-b | 0.0277 | 0.1376 | 0.1535 |
| VN-a - DMN-a | <0.0001 | 0.0021 | 0.0019 | DMN-b - DAN-a | <0.0001 | 0.0054 | <0.0001 |
| VN-a - DMN-b | <0.0001 | <0.0001 | <0.0001 | DMN-b - DAN-b | <0.0001 | 0.0054 | <0.0001 |
| VN-a - DMN-c | <0.0001 | <0.0001 | <0.0001 | DMN-b - VAN-a | 0.0011 | 0.0689 | 0.0019 |
| VN-a - TPN | <0.0001 | 0.0021 | <0.0001 | DMN-b - VAN-b | <0.0001 | 0.0021 | 0.0154 |
| VN-a - SN | <0.0001 | <0.0001 | <0.0001 | DMN-b - LN-b | 0.0238 | 0.1674 | 0.1059 |
| VN-b | <0.0001 | 0.0054 | 0.0176 | DMN-b - LN-a | <0.0001 | 0.0046 | 0.0216 |
| VN-b - SMN-a | <0.0001 | 0.0054 | 0.0588 | DMN-b - FPN-a | <0.0001 | 0.0054 | <0.0001 |
| VN-b - SMN-b | <0.0001 | 0.0046 | <0.0001 | DMN-b - FPN-b | <0.0001 | 0.0054 | 0.0166 |
| VN-b - DAN-a | <0.0001 | <0.0001 | <0.0001 | DMN-b - FPN-c | <0.0001 | <0.0001 | 0.0123 |
| VN-b - DAN-b | <0.0001 | <0.0001 | <0.0001 | DMN-b | <0.0001 | 0.0034 | 0.0019 |
| VN-b - VAN-a | <0.0001 | 0.0034 | 0.0145 | DMN-b - DMN-c | <0.0001 | 0.0034 | 0.0019 |
| VN-b - VAN-b | <0.0001 | 0.0021 | 0.1184 | DMN-b - TPN | <0.0001 | 0.0092 | 0.0300 |
| VN-b - LN-b | <0.0001 | 0.0034 | 0.0019 | DMN-b - SN | <0.0001 | <0.0001 | <0.0001 |
| VN-b - LN-a | <0.0001 | 0.0021 | 0.0123 | DMN-c - SMN-a | 0.0519 | 0.1901 | 0.1632 |
| VN-b - FPN-a | <0.0001 | 0.0021 | 0.0019 | DMN-c - SMN-b | 0.0142 | 0.0447 | 0.2105 |
| VN-b - FPN-b | <0.0001 | <0.0001 | 0.0206 | DMN-c - DAN-a | 0.0011 | 0.0327 | 0.0352 |
| VN-b - FPN-c | <0.0001 | 0.0067 | 0.0501 | DMN-c - DAN-b | <0.0001 | 0.0822 | 0.0087 |
| VN-b - DMN-a | <0.0001 | <0.0001 | 0.0019 | DMN-c - VAN-a | <0.0001 | <0.0001 | 0.0073 |
| VN-b - DMN-b | <0.0001 | <0.0001 | <0.0001 | DMN-c - VAN-b | 0.4182 | 0.4300 | 0.5600 |
| VN-b - DMN-c | <0.0001 | <0.0001 | 0.0019 | DMN-c - LN-b | <0.0001 | 0.0258 | <0.0001 |
| VN-b - TPN | <0.0001 | 0.0046 | 0.0019 | DMN-c - LN-a | <0.0001 | 0.0034 | 0.0087 |
| VN-b - SN | <0.0001 | 0.0152 | 0.0123 | DMN-c - FPN-a | <0.0001 | 0.0046 | <0.0001 |
| DMN-a - SMN-a | 0.0034 | 0.0537 | 0.0800 | DMN-c - FPN-b | 0.0316 | 0.0541 | 0.2119 |
| DMN-a - SMN-b | <0.0001 | 0.0116 | 0.0345 | DMN-c - FPN-c | 0.0913 | 0.2513 | 0.2277 |
| DMN-a - DAN-a | <0.0001 | 0.0021 | 0.0019 | DMN-c | 0.0044 | 0.0712 | 0.1100 |
| DMN-a - DAN-b | <0.0001 | <0.0001 | <0.0001 | DMN-c - TPN | <0.0001 | 0.0021 | 0.0117 |
| DMN-a - VAN-a | <0.0001 | <0.0001 | <0.0001 | DMN-c - SN | 0.6340 | 0.2653 | 0.8680 |

Table S-7. The obtained p-values of the emotion prediction results after FDR correction at the later stimulation stage. The highlights are the intra- and inter-networks with a significant prediction of both happiness and sadness ($p<0.0001$, FDR corrected).

| **Networks** | **Overall** | **Happiness** | **Sadness** | **Networks** | **Overall** | **Happiness** | **Sadness** |
|---|---|---|---|---|---|---|---|
| VN-a | <0.0001 | 0.0069 | <0.0001 | DMN-a - VAN-b | <0.0001 | <0.0001 | <0.0001 |
| VN-a - VN-b | <0.0001 | <0.0001 | <0.0001 | DMN-a - LN-b | <0.0001 | 0.0030 | 0.0339 |
| VN-a - SMN-a | <0.0001 | 0.0533 | 0.0025 | DMN-a - LN-a | 0.0122 | 0.0236 | 0.2088 |
| VN-a - SMN-b | <0.0001 | 0.0050 | 0.0243 | DMN-a - FPN-a | <0.0001 | 0.0030 | 0.0812 |
| VN-a - DAN-a | <0.0001 | 0.0050 | <0.0001 | DMN-a - FPN-b | <0.0001 | <0.0001 | 0.0025 |
| VN-a - DAN-b | <0.0001 | 0.0204 | <0.0001 | DMN-a - FPN-c | <0.0001 | 0.0136 | 0.0089 |
| VN-a - VAN-a | <0.0001 | <0.0001 | 0.0025 | DMN-a | <0.0001 | <0.0001 | 0.0057 |
| VN-a - VAN-b | <0.0001 | <0.0001 | <0.0001 | DMN-a - DMN-b | <0.0001 | <0.0001 | <0.0001 |
| VN-a - LN-b | <0.0001 | 0.0114 | 0.0057 | DMN-a - DMN-c | <0.0001 | <0.0001 | 0.0472 |
| VN-a - LN-a | <0.0001 | 0.0227 | <0.0001 | DMN-a - TPN | <0.0001 | <0.0001 | 0.0104 |
| VN-a - FPN-a | <0.0001 | 0.0069 | <0.0001 | DMN-a - SN | <0.0001 | 0.0258 | 0.0196 |
| VN-a - FPN-b | <0.0001 | <0.0001 | <0.0001 | DMN-b - SMN-a | 0.0229 | 0.2523 | 0.0559 |
| VN-a - FPN-c | <0.0001 | <0.0001 | <0.0001 | DMN-b - SMN-b | <0.0001 | 0.0207 | 0.0025 |
| VN-a - DMN-a | <0.0001 | 0.0147 | <0.0001 | DMN-b - DAN-a | <0.0001 | 0.0050 | <0.0001 |
| VN-a - DMN-b | <0.0001 | 0.0173 | <0.0001 | DMN-b - DAN-b | <0.0001 | 0.0050 | 0.0057 |
| VN-a - DMN-c | <0.0001 | 0.0084 | 0.0044 | DMN-b - VAN-a | 0.0013 | 0.0160 | 0.0967 |
| VN-a - TPN | <0.0001 | 0.0217 | <0.0001 | DMN-b - VAN-b | <0.0001 | 0.0098 | <0.0001 |
| VN-a - SN | <0.0001 | 0.0448 | <0.0001 | DMN-b - LN-b | <0.0001 | 0.0147 | 0.0563 |
| VN-b | <0.0001 | 0.0207 | 0.0057 | DMN-b - LN-a | 0.0024 | 0.1547 | 0.0542 |
| VN-b - SMN-a | 0.0101 | 0.0529 | 0.0694 | DMN-b - FPN-a | <0.0001 | 0.0050 | 0.0427 |
| VN-b - SMN-b | 0.0046 | 0.1905 | 0.0669 | DMN-b - FPN-b | <0.0001 | 0.0033 | 0.0147 |
| VN-b - DAN-a | <0.0001 | <0.0001 | <0.0001 | DMN-b - FPN-c | <0.0001 | <0.0001 | 0.0025 |
| VN-b - DAN-b | <0.0001 | 0.0098 | <0.0001 | DMN-b | <0.0001 | 0.0050 | 0.0147 |
| VN-b - VAN-a | 0.0024 | 0.0777 | 0.0160 | DMN-b - DMN-c | <0.0001 | 0.0443 | 0.0253 |
| VN-b - VAN-b | <0.0001 | 0.0069 | 0.0057 | DMN-b - TPN | <0.0001 | <0.0001 | 0.0147 |
| VN-b - LN-b | <0.0001 | 0.0136 | 0.0233 | DMN-b - SN | <0.0001 | 0.1085 | 0.0044 |
| VN-b - LN-a | 0.0024 | 0.0181 | 0.1315 | DMN-c - SMN-a | 0.0221 | 0.0084 | 0.4570 |
| VN-b - FPN-a | <0.0001 | <0.0001 | <0.0001 | DMN-c - SMN-b | 0.0024 | 0.1943 | 0.0222 |
| VN-b - FPN-b | <0.0001 | <0.0001 | <0.0001 | DMN-c - DAN-a | <0.0001 | 0.0207 | 0.0073 |
| VN-b - FPN-c | <0.0001 | 0.0098 | 0.0025 | DMN-c - DAN-b | <0.0001 | 0.0878 | <0.0001 |
| VN-b - DMN-a | <0.0001 | <0.0001 | <0.0001 | DMN-c - VAN-a | 0.0213 | 0.0388 | 0.1968 |
| VN-b - DMN-b | <0.0001 | 0.0181 | 0.0057 | DMN-c - VAN-b | <0.0001 | 0.0136 | 0.0603 |
| VN-b - DMN-c | <0.0001 | 0.0537 | 0.0276 | DMN-c - LN-b | 0.0677 | 0.0388 | 0.4203 |
| VN-b - TPN | <0.0001 | <0.0001 | <0.0001 | DMN-c - LN-a | 0.0141 | 0.0513 | 0.1899 |
| VN-b - SN | 0.1080 | 0.5060 | 0.0967 | DMN-c - FPN-a | <0.0001 | 0.0033 | 0.0025 |
| DMN-a - SMN-a | <0.0001 | 0.0084 | 0.0185 | DMN-c - FPN-b | <0.0001 | 0.0136 | 0.0044 |
| DMN-a - SMN-b | <0.0001 | 0.0193 | 0.0173 | DMN-c - FPN-c | <0.0001 | 0.0050 | 0.0606 |
| DMN-a - DAN-a | <0.0001 | 0.0033 | 0.0025 | DMN-c | 0.0069 | 0.0136 | 0.2718 |
| DMN-a - DAN-b | <0.0001 | 0.0050 | 0.0044 | DMN-c - TPN | 0.0141 | 0.1243 | 0.0967 |
| DMN-a - VAN-a | <0.0001 | <0.0001 | 0.0073 | DMN-c - SN | 0.1003 | 0.4861 | 0.1459 |
